# Age-associated growth control modifies leaf proximodistal symmetry and enables leaf shape diversification

**DOI:** 10.1101/2024.04.02.587754

**Authors:** Xin-Min Li, Hannah Jenke, Sören Strauss, Yi Wang, Neha Bhatia, Daniel Kierzkowski, Rena Lymbouridou, Peter Huijser, Richard S. Smith, Adam Runions, Miltos Tsiantis

## Abstract

Biological shape diversity is often manifested in modulation of organ symmetry and modification of the patterned elaboration of repeated shape elements.^1–5^ Whether and how these two aspects of shape determination are coordinately regulated is unclear.^5–7^ Plant leaves provide an attractive system to investigate this problem, because they usually show asymmetries along their proximodistal axis, along which they can also produce repeated marginal outgrowths such as serrations or leaflets.^1^ One case of leaf shape diversity is heteroblasty, where the leaf form in a single genotype is modified with progressive plant age.^8–11^ In *Arabidopsis thaliana,* a plant with simple leaves, SQUAMOSA PROMOTER BINDING PROTEIN-LIKE9 (SPL9) controls heteroblasty by activating *CyclinD3* expression, thereby sustaining proliferative growth and retarding differentiation in adult leaves.^12^ However, the precise significance of SPL9 action for leaf symmetry and marginal patterning is unknown. By combining genetics, quantitative shape analyses, and time-lapse imaging, we show that, in *A. thaliana*, proximodistal symmetry of the leaf blade decreases in response to an age-dependent *SPL9* expression gradient, and that SPL9 action coordinately regulates the distribution and shape of marginal serrations and overall leaf form. Using comparative analyses, we demonstrate that heteroblastic growth reprogramming in *Cardamine hirsuta,* a complex-leafed relative of *A. thaliana,* also involves prolonging the duration of cell proliferation and delaying differentiation. We further provide evidence that SPL9 enables species-specific action of homeobox genes that promote leaf complexity. In conclusion, we identify an age-dependent layer of organ proximodistal growth regulation that modulates leaf symmetry and has enabled leaf shape diversification.

**In Brief:** Age-dependent regulation of proliferative growth and differentiation along the proximodistal axis of leaves modulates organ symmetry and marginal patterning in heteroblasty, and enabled the action of homeobox genes in complex leaf shape evolution.

**Highlights:**

- An age-dependent growth gradient underpins regulation of *A. thaliana* leaf symmetry
- SPL9-mediated heteroblastic growth control potentiates leaf histogenic capacity
- A common growth framework underlies heteroblasty in complex and simple leaves
- SPL9 enables the action of homeobox genes that promote leaf complexity

## RESULTS

### Age-dependent changes in proximodistal symmetry of the leaf blade in *A. thaliana*

A notable aspect of leaf development in *A. thaliana* is the basipetal progression of differentiation.^13–17^ How this genetically-controlled gradient affects leaf symmetry along the proximodistal (PD) axis has not yet been investigated. In *A. thaliana*, juvenile leaves show a round, symmetrical blade, whereas adult blades are more elongated particularly in the proximal part and bear prominent marginal protrusions called serrations (Figures 1A and S1B). By quantifying the symmetry of the blade outline, we found that adult blades are more asymmetrical than juvenile blades along the PD axis (Figure 1A). Notably, in mature adult leaves, distal blades are round and have only a few, non-prominent serrations, which can both be considered juvenile features. In contrast, the proximal blades of adult leaves show increased eccentricity and numbers of serrations, which can be considered to be more adult characteristics (Figures 1A and S1B).

**Figure 1.**
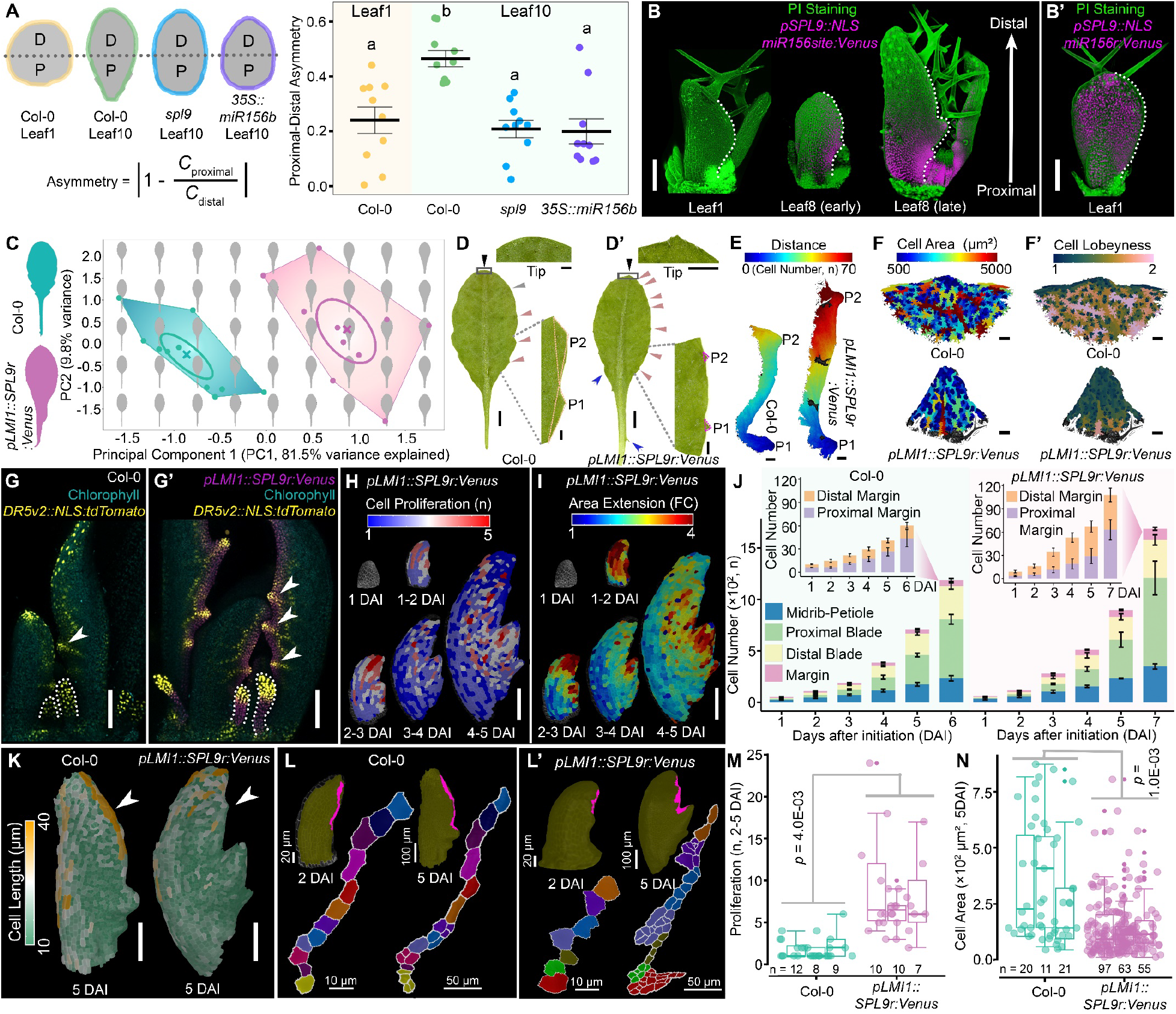
Distal *SPL9* augmentation disrupts proximodistal asymmetry in *Arabidopsis thaliana* adult leaf development. (A) Proximodistal asymmetry of leaf blade shape. *C*, curvature of the blade outline (see STAR Methods); D, distal; P, proximal. a and b, one-way ANOVA followed by posthoc Tukey’s Honest Significant Difference (HSD) test. (B) *SPL9* expression. Confocal projections of *SPL9* transcriptional reporters with (B) or without (B’) the regulation of miR156. Cells are outlined by propidium iodide (PI) staining. (C) Morphometric changes by *pLMI1::SPL9r:Venus.* Leaf10. Cross, mean; ellipse, SE; polygon, distribution range. (D and D’) Serration distribution and geometry (P1, P2; the first two proximal serrations, outlined by triangles) in representative leaf10. Triangles pinpoint serrations (gray triangle, serration relic along the distal margin in Col-0; black triangles, tip serrations). Blue arrowheads, stipule overgrowth. n > 10. (E-F’) Serration spacing (E, S1-S2) and maturation (F, cell area; F’, lobeyness. Tip serration, 600 μm radius) in mature leaf8. (G and G’) Serration development (white arrowheads, auxin maxima by *DR5v2*) in leaf8. Stipules, outlined. n > 5. (H and I) Heatmaps of cell proliferation (H) and area extension (I) for *pLMI1::SPL9r:Venus* leaf8. (J) Cell number increase in various cell types. Insets, marginal cells highlighted with proximal/distal classification. Error bars, mean ± SE. (K) Cell length heatmaps. Arrowheads, significant difference along distal margins. (L-N) Sector tracing of distal margin (in magenta) growth in Col-0 (L) and *pLMI1::SPL9r:Venus* (L’) leaf8. Cell proliferation (M) and cell area reached (N) in these sectors were quantified. Error bars, mean ± SE. *p*-value, nested ANOVA. Scale bars: 100 μm (B, B’, F-I and K); 200 μm (E); 5 mm, whole leaves and 1 mm, serrations in (D and D’); or indicated (K and L). n = 10 (A-D’), ≥ 3 biological replicates (E-N). DAI, days after initiation (H-N). See also Figure S1 and Video S1.

Recently, we found that SPL9 modulates heteroblastic changes in leaf shape by retarding differentiation and retaining growth along an age-dependent PD gradient, through the maintenance of proliferation by CYCD3.^12^ Given the basipetal decline in *SPL9* expression during adult leaf development (Figure 1B and Li et al.^12^), it is possible that SPL9-mediated control of growth contributes to the PD asymmetry of adult blades. To investigate this possibility, we first surveyed the changes in blade PD symmetry in *A. thaliana* genotypes with compromised SPL9 function. We found that PD asymmetry in adult blades is reduced in *spl9* and *miR156*-overexpressing (the microRNA miR156 negatively regulates *SPL9* expression^18,19^) plants (Figure 1A). These findings suggest that SPL9 is required to increase PD asymmetry in adult leaves.

### Distal augmentation of *SPL9* rebuilds proximodistal symmetry in adult leaves

To test whether SPL9 is sufficient to change the PD symmetry of leaf blades, we perturbed the PD asymmetry of *SPL9* expression in *A. thaliana* adult leaves. To this end, we ectopically augmented *SPL9* distal expression, in the context of its endogenous PD expression gradient (Figure 1B), using the regulatory sequences of *LMI1*, which predominantly confer expression in the distal margin of the leaf blade, in addition to stipules (Figure S1J and Vuolo et. al^20^).

We found that the activity of *pLMI1::SPL9r:Venus* (*SPL9r*, miR156-*r*esistant *SPL9* allele^21–23^) disrupts three different aspects of adult leaf asymmetry along the PD axis. First, it converts the asymmetrical adult blade to a more symmetrical oval shape (Figures 1C-1D’ and S1A). Second, it promotes PD homogeneity of serration distribution along the leaf margin (Figures 1D, 1D’ and S1B) by promoting serration establishment, especially in distal margins (Figures 1D, 1D’, 1G, 1G’, S1B, S1I and S1I’). This likely results from SPL9-dependent proliferation maintenance because more and smaller cells are present between serrations in *pLMI1::SPL9r:Venus* leaves than in wild-type (Figures 1E and S1D-S1F). Third, it causes serrations to be more symmetrical and uniform in size (Figures 1D, 1D’, and S1C), by inhibiting cell expansion and differentiation (Figures 1F, 1F’, S1G and S1H). In conclusion, forcing *SPL9* expression to be more uniform along the leaf margin increases the symmetry of both the leaf blade and individual serrations, and makes the organ-level distribution of serrations more uniform.

Next, we used time-lapse imaging to analyze the cellular growth pattern that confers PD symmetry to the *pLMI1::SPL9r:Venus* leaf8 (Video S1). We found that, compared to wild-type leaf8,^12^ the growth pattern was reprogrammed such that proliferative growth was prolonged and cell expansion was constrained, especially in the distal blade (Figures 1H-1K and S1K-S1Q). This reprogramming promotes marginal serration development (Figure 1K), and fate-mapping showed that it increases the contribution of distal cells to leaf development (Figures S1R-S1T). We also tracked the numbers of cells in distinct domains during leaf development, and found that these were boosted by the ectopic expression of *SPL9*, especially in margins (Figure 1J). Correspondingly, the PD distribution of cell proliferation along the margin, which shows more proliferation in the proximal part than distal part in wild-type, was evened out by *pLMI1::SPL9r:Venus* expression (Figure 1J), consistent with the increased number and uniform distribution of serrations observed (Figures S1B, and S1I’). By tracing the *pLMI1::SPL9r:Venus*-expressing sector in the leaf distal margin (Figure 1L’), we found that these cells grew at a similar rate (Figure S1U) as their wild-type equivalents (Figure 1L), but yielded more and smaller cells (Figures 1M and 1N), confirming that SPL9 is sufficient to protect cell proliferation from rapid differentiation and elongation at leaf margins. Altogether, these observations indicate that the basipetal decline of SPL9-dependent cell proliferation contributes to the establishment of morphological asymmetry along the PD axis in adult leaves.

### Proximal expression of *SPL9* is sufficient to convert juvenile leaves to adult shape

Considering that *SPL9* expression increases with leaf node progression and decreases along the PD axis during leaf maturation (Figure 1B and Li et al.^12^), we reasoned that the proximal expression of *SPL9*, which emerges through adult leaf development, might be critical for the progression to PD asymmetry. To directly test this idea, we evaluated whether expressing *SPL9r* proximally using the *CUC2*^24^ and *RCO*^20^ promoters (Figures 2B and 2B’) is sufficient to cause PD asymmetry in Col-0 juvenile leaves that lack native *SPL9* expression. In both genotypes, round juvenile (leaf1) blades were converted to an asymmetrical adult shape with an elongated base, similar to that observed in *pSPL9::SPL9r:Venus* (Figure 2A). This highlights the morphogenetic importance of local (proximal) SPL9 action in regulating heteroblastic changes in leaf symmetry.

**Figure 2.**
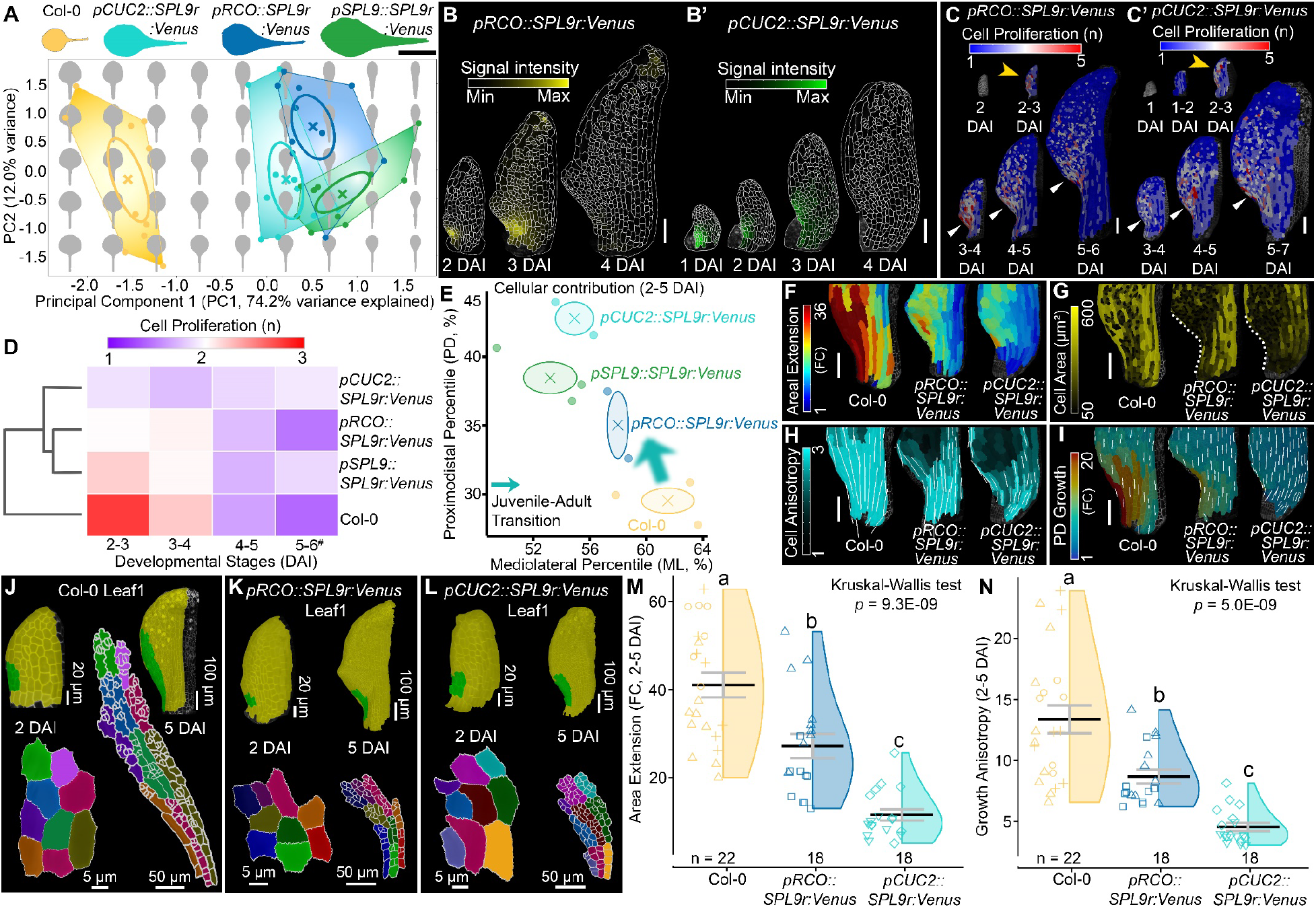
Proximal expression of *SPL9* is sufficient for juvenile to adult leaf shape transformation in *Arabidopsis thaliana*. (A) Shape-space plot of leaf1 with different genotypes. Cross, mean; ellipse, SE; polygons, distribution range. n = 10. (B and B’) *pRCO::SPL9r:Venus* (B) and *pCUC2::SPL9r:Venus* (B’) epidermal expression. (C and C’) Proliferation heatmaps for *pRCO::SPL9r:Venus* and *pCUC2::SPL9r:Venus* leaf1. Golden arrowheads, loss of proliferation burst; triangles, prolonged proliferation in proximal blades. (D) Hierarchical clustering of proliferation temporal dynamic. ^#^, 5-7 DAI for *pCUC2::SPL9r:Venus* and *pSPL9::SPL9r:Venus*. (E) Spatial shifts of cellular contribution by proximal SPL9. Cross, mean; ellipse, standard deviation. (F-I) Heatmaps of area extension (F), cell area (G), anisotropy (H), and proximodistal (PD) growth (I) in proximal blade-petiole regions. Measures, 2-5 DAI (cell area, 5 DAI), visualized on 5 DAI. White lines: growth direction where anisotropy > 1.5 in (H), PD direction in (I). (J-N) Growth tracing of ectopic *SPL9*-sectors. Col-0 equivalents (J) and *SPL9* highly-expressing cells in *pRCO::SPL9r:Venus* (K) and *pCUC2::SPL9r:Venus* (L) were traced from 2 to 5 DAI. Area extension (M) and anisotropy (N) were plotted, with biological replicates indicated in different shapes. Error bars, mean ± SE. Lowercase letters, post hoc Wilcoxon (FDR). Two (*pRCO::SPL9r:Venus* and *pCUC2::SPL9r:Venus*) or three (*pSPL9::SPL9r:Venus* and Col-0)^12^ biological replicates were analyzed (B-N). Scale bars: 1 cm (A); 100 μm (B-C’, and F-I); or indicated (J-L). DAI, days after initiation (B-N). See also Figure S2 and Video S2.

To then determine how cellular growth is altered by this proximal *SPL9* expression, we examined cell growth patterns in *pCUC2::SPL9r:Venus* and *pRCO::SPL9r:Venus* leaf1 using time-lapse imaging (Figure S2 and Video S2). We found that cell proliferation was prolonged at the base of leaf primordia, and that the “proliferation burst” that characterizes juvenile growth patterns^12^ disappeared in both transgenic lines (Figures 2C-2D and S2H). Correspondingly, proximal expression of *SPL9* reprograms the juvenile leaf proliferation pattern towards an adult one (Figure 2D, 2E and S2I-S2J”). Concomitantly, growth rate, anisotropy, and cell expansion and maturation are suppressed in *pCUC2::SPL9r:Venus* and *pRCO::SPL9r:Venus* leaves (Figures S2A-S2Gs), especially in proximal regions (Figures 2F-2I and S2K-S2N). To gain more insight into growth reprogramming upon local *SPL9* expression in juvenile leaves, we tracked sectors of *SPL9*-expressing cells and their wild-type equivalents (Figures 2J-2L). Compared to wild-type sectors that experience a proliferation burst, both *pCUC2::SPL9r:Venus* and *pRCO::SPL9r:Venus* sectors show repressed growth rate and anisotropy (Figures 2M and 2N), thereby retarding the elongation of the petiole and its delimitation from the blade (Figures S2A-S2E and S2O-S2S). These findings are in line with the action of SPL9 in heteroblastic reprogramming,^12^ whereby SPL9 slows proliferative growth and anisotropy while prolonging their duration, highlighting a temporal trade-off in the regulation of these growth parameters.

In conclusion, our results indicate that the persistence of *SPL9* expression in basal regions of adult leaves, in the context of its basipetal decline during primordia development (Figure 1B and Li et al.^12^), plays a fundamental role in breaking the symmetry of the leaf blade along the PD axis during heteroblasty. In this way, SPL9, with temporal inputs from both inter-leaf (increase with age) and intra-leaf (basipetal decline with maturation) levels, coordinately regulates proliferation competence and growth/differentiation patterns to modulate developmental symmetry.

### Evolutionarily conserved heteroblastic growth reprogramming enhances morphogenetic capacity in simple and complex leaves

In addition to symmetry modulation, expression of *pLMI1::SPL9r:Venus* along leaf margins results in increased serration number (Figures 1D’, S1B and S1I’), indicating that SPL9-mediated heteroblastic growth regulation is integrated with the marginal patterning system^17,24,25^ and influences the morphogenetic capacity of the leaf margin. Consistent with this idea, we found a positive correlation between SPL9 function and marginal serration development in *A. thaliana* leaves with perturbed *SPL9* expression (*SPL9* loss- and gain-of-function and *miR156* up- and down-regulated genetic lines, Figures 3A and 3B).

**Figure 3.**
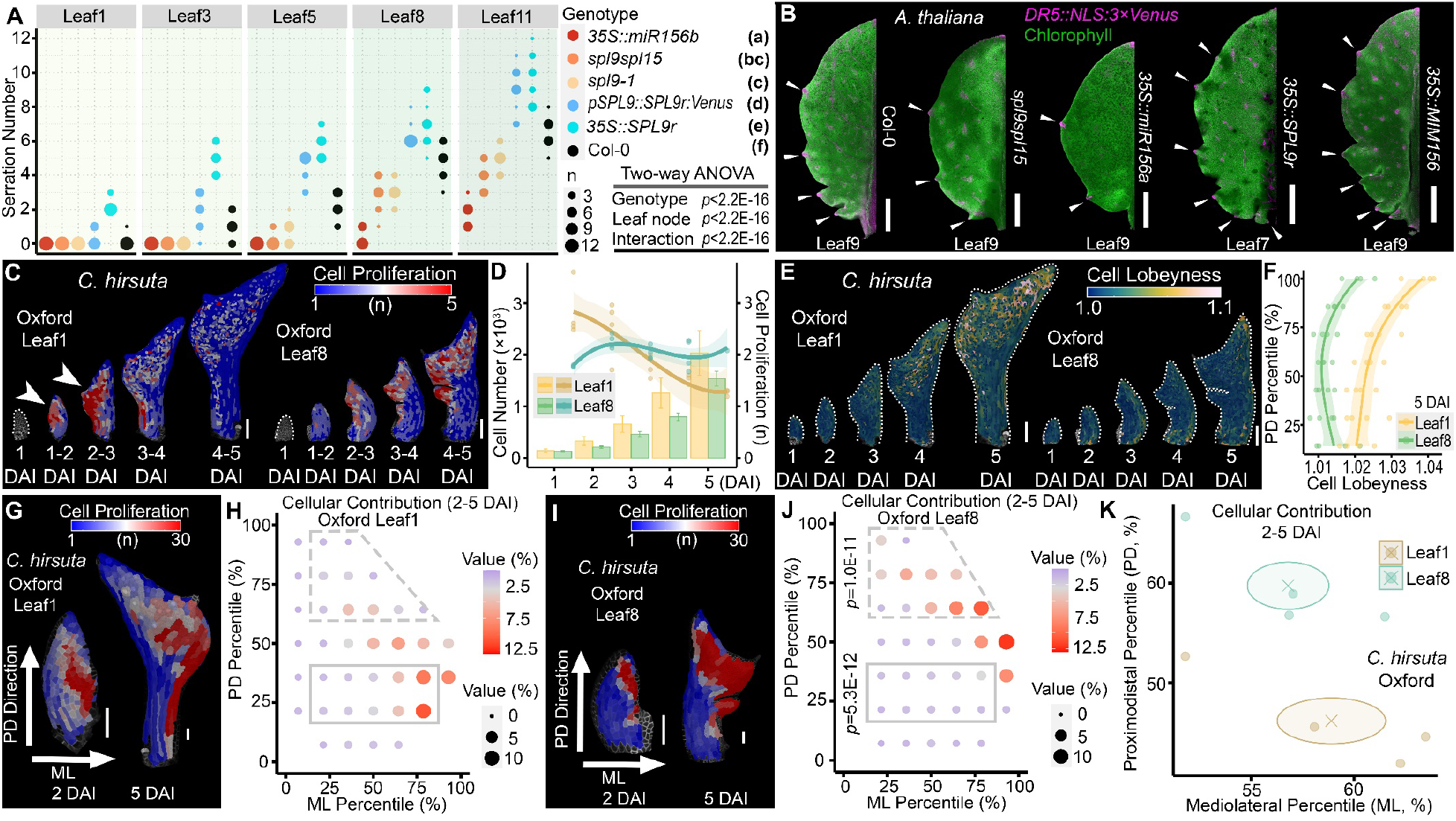
SPL9 facilitates serration development in *Arabidopsis thaliana*, and cellular growth reprogramming underlying heteroblasty in *Cardamine hirsuta*. (A and B) SPL9 promotes serration development in *A. thaliana*. (A), serration numbers in mature leaves. N = 12. Lowercase letters in the brackets, ANCOVA followed by post-hoc Tukey HSD. (B), serration establishment marked by *DR5* (triangles). Representatives, n > 6. (C and D) Spatiotemporal pattern of proliferation in *C. hirsuta* leaf1 versus leaf8. (C), proliferation heatmaps. Arrowhead, proliferation burst. (D), temporal dynamics of proliferation (right *y*-axis, dots with fitting lines) and consequent cell number accumulation (left *y*-axis, bars with mean ± SE). (E and F) Cell maturation in *C. hirsuta* leaf development. (E), heatmaps of cell lobeyness. (F), proximodistal (PD) alignment of cell lobeyness at 5 days after initiation (DAI). (G-K) Heteroblastic redistribution of cell proliferation, thus its developmental contribution, from 2-5 DAI. Proliferation from 2-5 DAI was projected on both 2 DAI (organ coordinates indicated; ML, mediolateral) and 5 DAI meshes (G, leaf1; I, leaf8). Cellular contribution was plotted in the two-dimensional coordinates (H and J), and the weighted centers were summarized (K). Ellipse, standard deviations of mean (inner cross) along PD and ML directions. The contribution of distal or proximal cells was compared between leaf1 (H) and leaf8 (J). *p*-values, two-way ANOVA. n = 4 biological replicates (C-K). Fitting lines, cubic regression with 95% confidence interval (D and F). Scale bars: 1 mm (B); 100 μm (C and E); 50 μm (G and I). See Figure S3 and Video S3.

The patterned formation of different types of marginal outgrowths, such as serrations, lobes, and leaflets, is a major source of leaf shape diversity in seed plants.^1^ Considering that the simple leaf of *A. thaliana* is a derived state that evolved via processes involving the acceleration of differentiation,^26^ we asked whether heteroblastic growth regulation, especially the proliferation burst associated with accelerated differentiation that occurs in *A. thaliana* juvenile leaf development,^12^ is part of these evolutionary processes. To address this question, we examined heteroblastic growth regulation in the *A. thaliana* complex-leafed relative *C. hirsuta* (leaf1 versus leaf8 live-imaging, Figure S3 and Video S3), which shows a heteroblastic increase in leaflet number, with no leaflets being present in leaf1^27^. In *C. hirsuta*, we found that cell proliferation (Figures 3C and 3D), along with areal growth (Figures S3A and S3A’), abruptly exhausts in a “burst” at the early stages of leaf1 development, while lasting longer in leaf8. This pattern of growth is similar to that which occurs in *A. thaliana*,^12^ thus indicating that the proliferation burst is a conserved and essential feature of juvenile leaves (in crucifer plants at least) and is unlikely to have contributed to *A. thaliana* leaf simplification in evolution. Correspondingly, cell expansion and differentiation are retarded in *C. hirsuta* leaf8 compared with leaf1 (Figures 3E, 3F, S3C-S3F’, and S3G), as is growth anisotropy at the leaf base (Figures S3B, S3B’ and S3G) that is associated with petiole differentiation. Fate-mapping revealed that proximal parts of leaf primordia contribute more to leaf development in leaf1 than in leaf8 (Figures 3G-3K and S3H-S3M), reflecting a delay in basipetal differentiation progression in leaf8 (Figures S3N and S3O), thus also mirroring comparisons seen in *A. thaliana* juvenile versus adult fate maps.^12^

### SPL9 as an enabler of leaf complexity evolution

The observed evolutionary conservation of heteroblastic growth reprogramming between *A. thaliana* and *C. hirsuta* raises the question of how it interacts with the developmental processes that distinguish simple from complex leaves. To address this question, we investigated whether heteroblastic reprogramming affects the action of two different types of homeobox genes, Class I *KNOTTED1-LIKE HOMEOBOX* (*KNOX1*) and HD-ZIP I *REDUCED COMPLEXITY* (*RCO*), which act in a species-specific manner to promote the generation of leaflets instead of serrations.^26,28–37^ To this end, we took advantage of the fact that *KNOX1* and *RCO* genes are sufficient to increase leaf complexity when ectopically expressed in *A. thaliana* leaves.^17,38,39^ We found that they create lobes or leaflets in adult (leaf8) but not juvenile leaves (leaf1) of *A. thaliana* (Figure 4A), despite discernible expression in both (Figures 4B-4D’) - which is also the case in the *C. hirsuta* endogenous context (Figures 4E-4F’). These observations indicate that the adult developmental context is an essential prerequisite for these two homeobox genes to increase leaf complexity.

**Figure 4.**
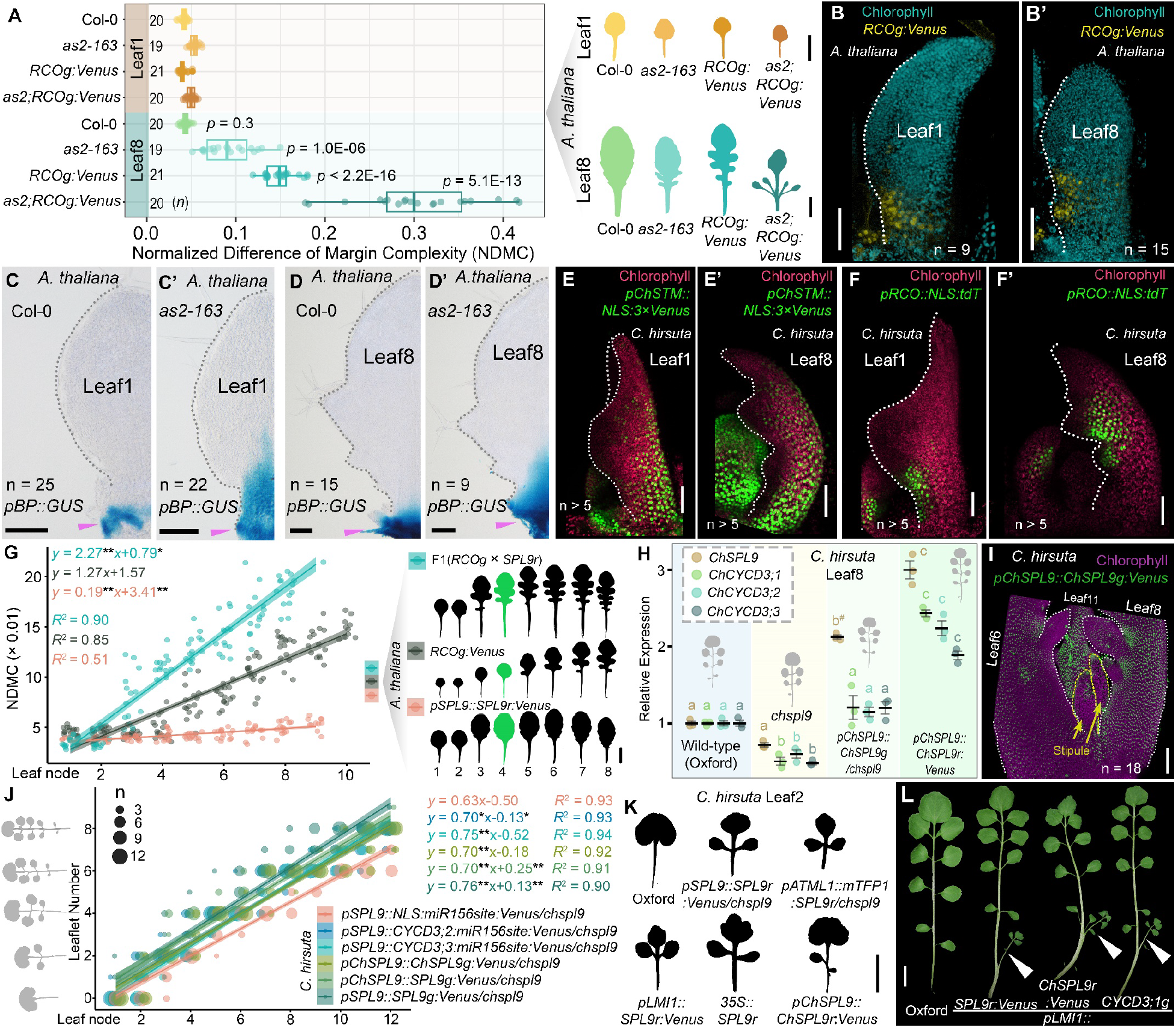
SPL9-sustained proliferation potentiates leaf morphogenesis in both simple and complex leaves. (A) Age-dependent modulation of *KNOX1* and *RCO* action in increasing *Arabidopsis thaliana* leaf complexity. *p*-value, Welch’s *t*-test (leaf1 versus leaf8) for each genotype. Right, representative leaf shapes in the plot colors. (B and B’) Heterologous expression of *RCO* at the lateral side of the bases of *A. thaliana* developing leaf1 and leaf8 (B’). (C-D’) *BREVIPEDICELLUS*/*KN1-like in Arabidopsis thaliana1* (*BP*/*KNAT1*), a *KNOX1* gene, does not express in Col-0 wild-type leaves (C and D), whereas, it expresses in the leaf base when the *KNOX* repressor AS2 is absent in both leaf1 (C’) and leaf8 (D’) of *as2-163* mutant.^39^ As a positive control, *BP* expression in axillary meristems is indicated (pink triangle). (E-F’) Confocal projections showing the expression of the homeobox genes *SHOOTMERISTEMLESS* (*STM*, E and E’) and *RCO* (F and F’) in *Cardamine hirsuta* juvenile (leaf1, E and F) and adult (leaf8, E’ and F’) leaves. (G) *SPL9* enables RCO to create lobes in *A. thaliana* leaves. n = 14. Right, genotype legend with representative silhouettes. Leaf4 in green highlights lobe emergence in the hybrid but not parents. (H) RT-qPCR examination. Lowercase letters, ANOVA-Tukey HSD with cut-off at 0.05 for each gene (indicated by colors). For *ChSPL9*, both transgenic and endogenous transcripts are included; ^#^, presence of non-functional *chspl9* transcripts. Error bars, mean ± SE; n = 3. (I) Confocal projection of *ChSPL9* expression in *C. hirsuta* adult leaves, with developing leaflets highlighted. n = 12. (K) Ectopic expression of *SPL9* is sufficient for precocious leaflet formation in *C. hirsuta* leaf2. n > 10. (L) *SPL9-CYCD3* in *LMI1* domain converts *C. hirsuta* stipule into an ectopic leaf (triangle, leaf12). n > 10. Linear model was established to show the age-dependent increase in leaf complexity (G and J), differences between control lines (G, *RCOg:Venus*; J, *pSPL9::NLS:miR156site:Venus/chspl9*) and indicated genotypes were tested by ANCOVA (slope and intercept); *, *p* < 0.05; **, *p* < 0.01. shade, 95% confidence interval. Scale bars, 100 μm (C-D’, and I); 50 μm (B, B’, and E-F’); 1 cm (A, G, K and L). See Figure S4.

Next, we showed that this developmental enabling effect by heteroblasty is miR156-SPL9 dependent, because *SPL9* overexpression (Figure 4G) or *miR156* knockdown (Figure S4A) strongly enhanced the ability of RCO to generate deep lobes in *A. thaliana* leaves, whereas *miR156* overexpression has the opposite effect (Figure S4A’). We then probed the morphogenetic role of SPL9-mediated heteroblastic reprogramming in leaflet formation in *C. hirsuta*. Like its ortholog in *A. thaliana*,^12^ *ChSPL9* expression, under the control of miR156, increases with shoot age and declines with leaf maturity (Figures S4B-S4C”). Interestingly, ChSPL9 accumulates at the base of developing leaflets (Figure 4I), a phenocritical domain where genetically-controlled growth and proliferation modulate leaflet production in *C. hirsuta*.^17,39,40^ Heteroblastic progression in *C. hirsuta* is delayed in genetic lines with compromised ChSPL9 function and accelerated when *ChSPL9* expression is increased (Figure S4D). In addition, *CyclinD3* family genes, which are major downstream effectors of SPL9 action in prolonging proliferative growth in *A. thaliana* heteroblasty,^12^ also show ChSPL9-dependent temporal expression in the development of *C. hirsuta* complex leaves (Figures 4H, S4B, and S4B’). This highlights that SPL9-CYCD3 is an evolutionarily conserved module in heteroblastic reprogramming. We also found that *A. thaliana SPL9* or *CYCD3* genes can rescue the *chspl9* heteroblastic defects to an equivalent level of the endogenous *ChSPL9* when expressed in the *SPL9* domain (Figure 4J), and that ectopic expression of *SPL9* from both species is sufficient to create leaflets in *C. hirsuta* juvenile leaves (Figure 4K). Moreover, expressing *SPL9* or *CYCD3;1* in the *LMI1* domain is sufficient to promote stipule overgrowth into an ectopic leaf in the adult leaf of *C. hirsuta* (Figure 4L), similar to that reported in *A. thaliana*.^12^

Overall, our data indicate that the SPL9-mediated growth reprogramming, associated with a differentiation delay, underlies heteroblasty in both simple and complex leaves and provides temporal input into distinct features of marginal outgrowths.

## DISCUSSION

Our work underscores the role of SPL9 as part of an age-dependent positional information system in leaf heteroblastic morphogenesis. This system tunes cellular growth along the PD axis, thereby influencing the PD symmetry of a leaf, mainly through prolonging proliferative growth and preventing differentiation. As such, organ-level regulation also provides a broader spatiotemporal window and more cells for the marginal patterning system^17,24^ to act upon and generate marginal protrusions. This growth-patterning interaction, in addition to previously studied protein-protein interactions,^25^ provides a cellular mechanism for SPL9-promoted marginal morphogenesis that is consistent with and therefore validates previous computational models of leaf development.^17,24^ Moreover, this heteroblastic growth reprogramming by SPL9 is evolutionarily conserved between *A. thaliana* simple leaves and *C. hirsuta* complex leaves, and encodes information that reads out as node-specific differences in leaf development along the PD axis – including blade shape, serration geometry and distribution in *A. thaliana*; and leaflet formation in *C. hirsuta*.

Our findings further indicate that a proliferation burst is a conserved feature of juvenile leaf primordia in both *A. thaliana* and *C. hirsuta*. Whereas the species-specific complex leaf shape regulators *RCO* and *KNOX* cannot exert their stimulatory effects on leaf complexity in such juvenile leaf developmental context, their effects become prominent in the context of SPL9-dependent heteroblastic growth reprogramming. This suppression of leaf complexity by the juvenile state can be viewed as a form of within-genotype developmental epistasis, where rapid growth and differentiation prevent the elaboration of the complex-leaf morphogenetic program. By the same token, the SPL9-mediated differentiation delay and growth reprogramming likely allowed the action of KNOX and RCO and consequent leaf shape diversification during evolution. Interestingly, the function of SPL9 in delaying differentiation can be considered comparable, although weaker, to that of KNOX1 proteins.^17,30,35,36^ This suggests that multilevel control of the progression of leaf differentiation provided evolution with the flexibility to explore a broader phenotypic space. Notably, natural allelic variation in *ChSPL9* underlies a heterochronic QTL (quantitative trait locus) that contributed to the evolution of a distinctive *C. hirsuta* morphotype prevalent in the Azorean islands.^41^ This QTL shows evidence for positive selection, and its distribution mirrors a climate gradient that broadly shaped the Azorean flora.^41^ Taken together, the effects of *SPL9* in leaf shape diversity, combined with genetic analyses of heteroblasty and its natural variation in different land plants including trees,^11,41–51^ indicate that the heterochronic regulation of growth that we report here constrained plant evolution at both the micro-and macro-evolutionary scales.

## ACKNOWLEDGMENTS

We thank Juliana Medina, Shuping Xing, and Claudia Canales for help with generating genetic materials, Eirini Tsaroucha and Liam McGuire for help with MGX analyses. We acknowledge R. Scott Poethig and Sarah Hake for sharing plant materials. We thank Angela Hay for critically reading the manuscript and Gemma Richards for manuscript production. This project was supported by a Max Planck Society core grant (M.T.), Bundesministerium für Bildung und Forschung (BMBF, Enhanced Crop Photosynthesis-2 031B0881B to M.T.), Deutsche Forschungsgemeinschaft (DFG, in terms of the multidisciplinary research unit FOR2581 “Quantitative Morphodynamics of Plants”, the Cluster of Excellence on Plant Sciences [CEPLAS], and the transregional collaborative research centre CRC TRR 341 “Plant Ecological genetics” to M.T.), and an Alexander von Humboldt Postdoctoral Fellowship to X.-M.L.

## AUTHOR CONTRIBUTIONS

X.-M.L. designed experiments with M.T., performed the majority of the work, and conducted data analyses and material validation. H.J., D.K., and R.L. were involved in time-lapse experiments, and S.S. and A.R. contributed to growth quantification and related methods. Y.W. performed the *as2;RCOg:Venus* assays. N.B. imaged *ChSTM* and *RCO* expression in *C. hirsuta*. P.H. and R.S.S. contributed to method development. X.-M.L. prepared the draft and figures. X.-M.L. and M.T. wrote the manuscript with input from other authors; M.T. conceived this project.

## EXPERIMENTAL MODEL AND SUBJECT DETAILS

### Plants and growth conditions

All genetic lines were created in the *Arabidopsis thaliana* Col-0 accession or *Cardamine hirsuta* Oxford accession.

Both species were cultivated under the same growth conditions after stratification at 4°C (3 days for *A. thaliana*; 5 days for *C. hirsuta*) as follows.

Soil-grown plants were cultivated in controlled climate chambers (light intensity: 120 µmol·m^-2^·s^-1^; temperature: 20°C/18°C for day/night; relative humidity, 60%). For leaf shape assays, plants were grown under short-day conditions (SD, 8-hr light / 16-hr dark) until target leaves were fully expanded, because these growth conditions can reduce leaf curling, making the leaves relatively flat and easy to measure. For time-lapse imaging, plants were grown under long-day conditions (LD, 16/8 hrs) until the target primordium initiated. Dissected plantlets were cultivated on ½ MS medium plates supplemented with 1% sucrose and 0.1% plant preservative mixture (PPM, Plant Cell Technology) for 24 hrs between two imaging events. For *RCO* and *STM* expression imaging in *C. hirsuta*, seeds were sterilized and grown on GM medium under LD conditions.^40^

To harvest seeds, (after genetic crosses, plant transformation and screening, or seed propagation) plants were grown in MPIPZ greenhouses under LD conditions. Basta screening was conducted on soil while hygromycin-resistance screen (25 µg/mL hygromycin and 300 μl/mL cefotaxime) was performed on ½ MS plates before transferring to soil.

## METHOD DETAILS

### Genetic materials

#### • Molecular cloning

Plasmids were constructed as follows (see Table S1 for primers used). Cloned fragments were validated by Sanger sequencing, and all plasmids were verified by Illumina sequencing before being used to transform plants via the floral dip method.^30,52^

For *SPL9* targeted expression, the SPL9r-Venus fragment was cloned from the *p*SPL9::SPL9r:Venus plasmid^12^ by PCR with NotIXmaI-SPL9-F/NotIXmaI-Venus-R, then cloned into the *p*RCO-pMLBart^20^ via *Not* I or subcloned into *p*LMI1-pBJ36^53^ and *p*CUC2-pBJ36^24^ via *Xma* I before *Not* I cloning into the destination vector pMLBart. For cloning using a single restriction enzyme (e.g., via *Xma* I or *Not* I), after digestion, calf intestinal alkaline phosphatase (CIP) was used to remove the phosphate groups from the 5’-termini of linearized vectors to prevent vector self-ligation.

For *ChSPL9* cloning, the NotI-gChSPL9:Venus-NotI and miR156-resistant version NotI-pChSPL9::ChSPL9r:Venus-NotI were synthesized (GenScript) and transferred to the pMLBart vector via *Not* I cloning. The *SPL9* promoter in construct pMDC32-*p*SPL9::SPL9:Venus and the SPL9r:Venus fragment in plasmid pMLBart-*p*LMI1::SPL9r:Venus were swapped for the *ChSPL9* promoter and ChSPL9r:Venus, to generate *p*ChSPL9::SPL9:Venus and *p*LMI1::ChSPL9r:Venus, respectively.

35S::SPL9r, 35S::miR156, and 35S::MIM156 were synthesized and cloned into pMLBart via *Not* I. The *p*ChSTM::NLS:3×Venus construct was made based on the previous *p*ChSTM::GUS reporter,^30^ with the β-glucuronidase coding sequence replaced by NLS: 3×Venus.

#### • Plant genetic stocks

Genetic crossing: fluorescent plasma membrane (*pUBQ10::PM:tdTomato*) or auxin activity reporter (*DR5::NLS:3×Venus*, or *DR5v2::NLS:tdTomato*) markers were introduced into relevant lines (representative transgenic lines or mutants) by crossing (indicated by ×), to facilitate imaging. The *RCOpro:RCOg-VENUS* (*RCOg:Venus*) transgenic line^20^ was crossed into genetic backgrounds with perturbed expression of *miR156-SPL9* (*pSPL9::SPL9r:Venus*, *35S::MIM156*, and *35S::miR156*).

The *A. thaliana as2* mutant, which exhibits elevated expression of *KNOX1* genes, was the *as2-163* allele obtained from a screening of an ethyl methanesulfonate (EMS) mutagenized library.^39^ It carries a loss-of-function allele of *AS2* with a point mutation (cytosine to thymine, leading to an amino acid substitution from Ala^65^ to Val).

The *C. hirsuta chspl9* mutant was created using CRISPR-Cas9 technology.^41^ It contains two single-base insertions within the first exon of *ChSPL9*, with a guanine (G) inserted after the 53^rd^ base of the coding sequence (CDS), causing a reading frame shift and a premature stop codon 15-bp afterwards, and a cytosine (C) inserted after the 179^th^ base of the CDS. A Cleaved Amplified Polymorphic Sequences (CAPS) marker was developed to genotype *chspl9* allele (see Table S1). Transgenic lines, mutants, and genetic combinations used in this study were genotyped using molecular markers (listed in Table S1, together with primers used for cloning). In addition to gel electrophoresis, PCR products were also validated by Sanger sequencing.

### Expression analysis

#### • Fluorescence imaging

We used a Leica TCS SP8 Confocal Laser Scanning Microscope (CLSM) for fluorescence imaging. Fluorescence signals were collected at channels as follows:

| Fluorescence | excitation wavelength | collection range |
| --- | --- | --- |
| Venus/YFP | 514 nm | 525-545 nm |
| tdTomato | 552 nm | 570-620 nm |
| PI (Propidium Iodide) | 552 nm | 610-640 nm |
| chlorophyll | 514 nm | 660-725 nm |

Leaf primordia or developing leaves were either imaged under water in growth plates with long working-distance water immersion objectives (AP 20×/0.75; AP 40×/1.10. e.g., 1-6 DAI samples in Figures S1I and S1I’, Figures 4E-4F’, 4I, and S4C-S4C”) or were mounted on slides and imaged using standard water objectives (AP 20×/0.80; AP 40×/0.50. e.g., 7-12 DAI samples in Figures S1I and S1I’ and Figure 3B). Imaging was performed at 200 Hz and 3-5 × line averaging and sections were spaced around 1 μm.

Expression of ectopic SPL9 in *pRCO::SPL9r:Venus* (Figure 2B), *pCUC2::SPL9r:Venus* (Figure 2B’), and *pLMI1::SPL9r:Venus* (Figure S1J) were imaged in parallel with time-lapse imaging of cell growth (as illustrated below) with the same settings within each group to facilitate comparisons. Epidermal (2-6 µm) Venus signals were projected onto the segmented cellular meshes in MorphoGraphX (MGX).^54,55^

#### • GUS staining

To examine the *pBP::GUS* expression in *A. thaliana* leaves, histochemical GUS detection assays were performed in both the *as2-163* mutant (Figures 4C’ and 4D’) and Col-0 wild-type (Figures 4C and 4D) backgrounds. Soil-grown plants were harvested at corresponding stages, and young shoots were fixed in cold 90% acetone (v/v) for 30 min. Then, samples were washed three times with staining buffer (2mM K4Fe(CN)6, 2mM K3Fe(CN)6, 10mM EDTA, and 0.1% Triton X-100 (v/v) in 50mM phosphate buffer, pH 7.2). For β-Glucuronidase (GUS) staining, samples were incubated in GUS staining solution (1 mM X-Gluc in the staining buffer) at 37°C for 10 hours after vacuum-infiltration. To stop the staining and to clean tissues, samples were then incubated in 20%, 50%, 70%, and 90% ethanol (v/v) sequentially, for 3 hrs in each solution with shaking. For imaging, samples were dissected under a stereomicroscope (Nikon SMZ1500) and imaged using a Nikon SMZ18 stereomicroscope.

#### • RT-qPCR

*C. hirsuta* developing leaves were harvested from soil-grown plants (LD conditions) at indicated stages. Three biological replicates were used for each test, with at least three leaves mixed for each replicate. Total RNA was extracted using the RNeasy Plant Mini Kit (QIAGEN) and first-strand cDNA was synthesized with the SuperScript® VILO^TM^ cDNA Synthesis Kit (Invitrogen). Real-time qPCR was carried out on a QuantStudio™ 3 Real-Time PCR System (Applied Biosystems) using Power SYBR^TM^ Green PCR Master Mix (Applied Biosystems) in a 20 μl reaction. Amplification efficiencies of qPCR primers (see Table S1) were assessed (95-100%) and relative quantification of target genes transcripts was performed using 2^-ΔΔCT^ method.

### Time-lapse growth study

#### • Live-imaging

Seedlings at corresponding stages were dissected under a stereomicroscope (Nikon SMZ1500) to expose the target primordia. They were then cultivated (LD) on square petri dishes with ½ MS medium and imaged on the same plates (submerged in water, which was removed afterwards) under long working-distance water immersion objectives (AP 20x/0.5 or AP 40x/0.8). Imaging was performed on a TCS-SP8 (Leica) upright confocal laser-scanning microscope. To minimize stressing tissues, confocal stacks were acquired with hybrid detectors (HyDs) at 512×512 resolution and a scanner speed of 400 Hz (bidirectional) without averaging. To optimize signal quality, imaging settings (e.g., the gain, pinhole, and sometimes the laser intensity) were adjusted among samples or during the imaging of the same sample. Sections were spaced from 0.5-1 μm apart in the *z*-dimension depending on the sample size. Large samples were scanned in part and overlapping stacks were stitched in MGX.^54,55^

#### • Growth measurement

Time-lapse images were analysed in the MGX software following the standard pipeline.^54,55^ Briefly, a curved-surface (2.5D) mesh was created and cells were segmented. Parental lineage between two time points was manually labelled and corrected after the “Check correspondence” examination, and lineages over various time windows were then combined with a custom python script (“combinelineages.r”^12^) to compute a full-lineage cell fate map.

Cell geometry, including cell area, length, lobeyness, and solidarity were measured. The degree of pavement cell undulation was estimated using cell lobeyness^56^ or solidarity. Cell solidarity (e.g., Figure S3E) is a ratio of the area of a cell’s convex hull over that of the cell. Similar to lobeyness, solidarity also reflects cell shape complexity, but is more robust to mesh noise compared to the former.

Cell correspondence was tracked over time (e.g., Figures 1L, 1L’, 2J-2L, S2R, S2S, S3H, and S3I-S3J’) by their parental lineages. Cellular growth (changes from a single cell at the start time point to the clonal sector including all of its daughter cells at the next time point), including proliferation (increase in cell number), area extension (relative increase in area), cell area change (the ratio between increases in area and in cell number),^12^ and growth anisotropy (the ratio between the amount of maximum and minimum Principal Directions of Growth, PDG)^54^ were computed using the “Growth Analysis 2D” process.

Organ coordinate axes along the proximodistal (PD) or mediolateral (ML) directions were established in Euclidean distance fields.^12,55^ A cell line at the bottom of a primordium or the midline of the primordium was manually selected as the reference position of the PD and ML axes, respectively. The Euclidean distances (shortest path) of each cell relative to these reference lines were calculated as its positional information in the organ coordinates.

Developmental contribution was calculated to evaluate the contribution of a cell group (a bin) to the total cell number (cellular contribution) or total area (areal contribution) of developing leaves after a specific period. It was calculated as the changes in the relative proportion of clones in the whole leaf after a period of growth, which was normalized by the initial cell number or area of the bin.^12^ For instance, areal contribution of a bin at 2 DAI to the 5 DAI leaf would be calculated in this way:

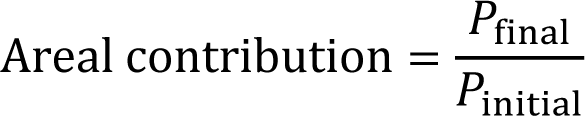

Where *P*initial is the proportion of the area of the target bin (total area of cells in the group) to the total area (only complete lineages over the measuring period were considered) of the 2 DAI primordium; *P*final is the proportion of the area of the tracked bin (total area of all the clones originating from the cells in the initial 2 DAI bin) to the total area (all the lineages tracked from 2-5 DAI) of 5 DAI leaf. The cellular contribution is computed with a similar formula, where area is replaced by cell number.^12^ Both relative cellular and areal contributions were transformed into percentages and plotted with biological replicates according to the positional information (bin-id). PD/ML growth was calculated as the relative increase (fold change, total length of the progeny clone / the length of the mother cell) in length along the PD and ML organ axes. Values were visualized on the start (e.g., Figures S1P and S1P’) or the end mesh (e.g., Figure S1L). For the latter, the coordinates were re-established using a custom Bezier grid at the target mesh.

Measures were exported to CSV files and further analyzed and plotted in R. Representative heatmaps are shown.

#### • Growth comparison

Organ-level growth was compared between diverse genotypes simply by the mean of cellular growth (e.g., the temporal changes in the average growth of leaves in Figure 2D). Growth was also compared at higher resolution, for instance, at sub-organ, tissue, or cell group level. Once cells have categorical identities, they can be grouped, which facilitates statistical analysis on cellular measures at the group level. These categorical factors can be the relative positions of cells in the organ coordinates, a fixed distance to a specific structure (e.g., the tip of a serration), or their developmental regions.^16,17,55^

### Developmental grouping

Given that different developmental domains show divergent growth patterns, it is informative to compare growth in distinct domains and among different genotypes. This was achieved based on cell fate maps. Considering that histogenic divergence increases over developmental time, we defined developmental domains at the latest time point according to the position, geometry, and growth features of cells (Figure S1Q). Specifically, the margin domain contains elongated giant cells at the edge of a leaf; the petiole-midrib domain is located along the central PD axis of the leaf and includes longitudinally elongated cells that grow relatively slowly; the blade domain is the region between the margin and the petiole/midrib and is composed of cells of variable sizes, shapes and growth rates. Leaf margin and blade can be further divided into distal and proximal parts according to the Euclidean distance along the PD axis (Figure 1J). These cell type identities were traced backwards in time according to cellular parental lineages. The average growth of different domains was tracked and analyzed (e.g., Figures 1J, S1Q, and S2H). Representative clones of different developmental paths are shown (e.g., Figures S2R, S2S, and S3I-S3J’).

### Expression domain

Developmental lineages were also defined by specific gene expression. In the targeted expression of *SPL9* experiments (*pLMI1::SPL9r:Venus, pRCO::SPL9r:Venus,* and *pCUC2::SPL9r:Venus*), to precisely monitor the local growth modulated by ectopic *SPL9* expression, we identified and tracked the ectopic-*SPL9* lineages. Sectors were defined based on transgene (*SPL9r:Venus*) expression and their developmental positions. For *pLMI1::SPL9r:Venus*, we focused on the distal margin (Figures 1L-1N) because it is the major expression domain of *LMI1*. It was defined as the marginal cell line from the leaf tip to the tip of the first serration (the largest serration) in both *pLMI1::SPL9r:Venus* and wild-type Col-0. In the SPL9 proximal supplementation study (Figure 2), a cell population consisting of nine cells was selected as the common domain of both *pRCO::SPL9r:Venus* and *pCUC2::SPL9r:Venus* at 2 days after initiation (DAI); the equivalent cell population in Col-0 at 2 DAI was identified according to the relative location (Figures 2J-2N). The growth of these target sectors was tracked from 2-5 DAI and compared with that of the control equivalents.

### Regional clustering

The distance field was determined to define a region of interest with key morphological features as landmarks. For example, to compare the cellular properties of serration development and spacing in Col-0 and *pLMI1::SPL9r:Venus* mature leaves, the distance between the first two proximal serrations was determined (Euclidean distance in Figure S1D and cell number in Figure 1E). Similar delimitation of a region was applied for the tip serration analysis in these two genotypes, where tip serration was defined with the tip cell as the center of a region with a radius of 600 µm (e.g., Figures 1F and 1F’). Cells in this sector were measured and compared between genotypes with biological replicates.

### Positional binning

Cellular growth from biological replicates and different genetic backgrounds was aligned to a common axis (e.g., the PD axis) or coordinates (PD-ML coordinates mentioned above) according to the relative positional information of cells.^16,55^ Growth alignment was performed at a bin level, where cells were grouped along a specific axis to get bins with equivalent cell number (cellular binning or ordinal binning), area (areal binning), or distance (Euclidean binning). The group information is then propagated by cell lineages across time points. The growth of each bin was calculated as the average growth of cells within the bin, and cells that were not captured throughout the growth tracing period were not included. Growth alignment was performed in either one-dimension (e.g., PD alignment graphs in Figures 3D and S3J-S3N) or two-dimensions (e.g., Figures 3H, 3J, and 3K). For PD alignment, ordinal binning was used; for 2D alignment, Euclidean binning was applied to make an effective binning with limited cells. The binning effect was afterwards removed by cell number or area normalization.

#### • Growth animation

Growth animation (Videos S1-S3) was created using deformation functions in MGX^55^, where cell lineages and corresponding landmarks were used to map one mesh onto the other, and the frames were thereby generated. Individual frames were then compiled in FFmpeg (https://ffmpeg.org) with titles, scale bars, and explanatory annotations incorporated.

### Quantitative morphometrics

#### • Blade shape asymmetry

The shape asymmetry of leaf blades along the PD axis was estimated by comparing the curvature of the blade outlines (Figure 1A). Fully expanded rosette leaves were collected and scanned. Leaf silhouettes were used for curvature analysis in Fiji. An open-source Fiji plugin, Kappa,^57^ was used for measuring curvature. Specifically, the outline of a blade was manually tracked by adding control points along its margin (from the tip to the blade-petiole junction) on one side. These control points were used by the software to build a corresponding B-splines curve using a least-squares based minimization algorithm. The fitting curve was checked and adjusted to make sure it matched the blade outline properly (serrations and sinuses were smoothed for the outline curvature estimation). Curvature was calculated in Kappa, together with PD position information. Exported data were analyzed in R; the asymmetry index was computed using this formula:

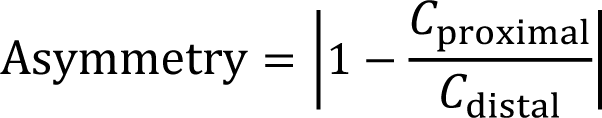

Where *C*proximal and *C*distal represent the average curvature of the proximal and distal blade, respectively.

#### • Leaf shape space

Digital silhouettes of mature leaves were quantitatively analyzed using a PCA-based multivariate shape analysis method using the Leaf Interrogator software (LeafI)^16^. Essentially, leaf contours were extracted from images; then two common landmarks (the tip of the blade and the base end of petiole) were specified for each contour to set a common axis for rotation invariant analysis. Leaf contour was resampled to obtain 120 points (semi-landmarks) between the two common landmarks. The resampled contours were used for shape geometry computation using translation and scale-invariant Elliptical Fourier Descriptor (EFD) method. Shape spaces were built by removing leaf bilateral asymmetry.

#### • Leaf complexity index

To quantify the marginal complexity introduced by the RCO/KNOX1 module in *A. thaliana*, we employed the Normalized Difference of Margin Complexity (NDMC) index,^58^ which represents a size-invariant measure of leaf dissection on a two-dimensional plane. NDMC was calculated as the perimeter of the leaf margin relative to its convex hull:

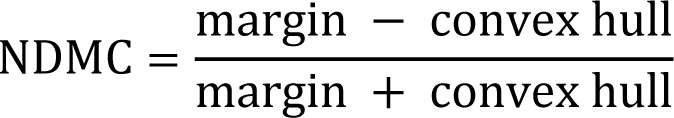

Where, the convex hull is the smallest polygon enveloping the whole leaf.

The NDMC was computed in LeafI software,^16^ and results were plotted in R (e.g., Figures 4A and 4G).

#### • Serration morphospace

To quantitatively investigate serration geometry in Col-0 and *pLMI1::SPL9r:Venus* mature leaves (Figure S1C), ten fully expanded leaf10s were scanned (as described above) for each genotype. Serrations were ordered from the tip following their developmental sequence. The first three serrations from both sides of each leaf were examined, except for the first in Col-0, which fades during leaf maturation and was thus unmeasurable. Intercalary serrations in *pLMI1::SPL9r:Venus* were also ignored, because there was no counterpart in the wild-type for comparison. Every serration was geometrically summarized as a triangle spanning the distal and proximal sinuses and terminating at the serration tip. The triangular frame of each serration was measured using FIJI, including the lengths of the three sides and the height. These triangular approximations were plotted in a morphospace with pairwise combinations of the width (basal distance between the distal and proximal sinuses), height (distance of the protrusion tip from the base) and asymmetry (difference between the proximal and distal sides of the triangle) as reported previously,^17^ while asymmetry was normalized by serration size (the height):

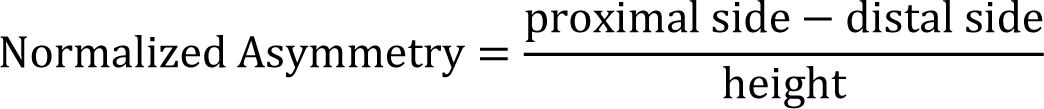

Here the distal versus proximal sinus/side refers to the leaf-level rather than the serration-level coordinate, e.g., the distal sinus is the junction between the serration and leaf edge towards the leaf tip, while the proximal sinus is the one closer to the leaf petiole. The prominence of serrations was captured by the height and width of the triangles, whereas the shape variance was depicted as the proximo-distal asymmetries.

Representative leaves (Figures 1D and 1D’) were imaged using a Zeiss Smartzoom 5 digital microscope.

## QUANTIFICATION AND STATISTICAL ANALYSIS

Quantitative data (e.g., growth data from MGX) were analyzed and visualized in R. All statistical tests and data size (*n*) have been indicated in figures or related legends. The *p*-value was provided whenever possible, otherwise the cut-off for statistical significance was 0.05. Parametric tests were preferably deployed once the assumptions were met; otherwise, non-parametric alternatives were used.

### • Parametric tests

1. for comparisons between two groups (e.g., Figure S1A), the student’s *t*-test was used;
2. for multiple comparisons (e.g., Figure 1A), one-way ANOVA (ANalysis Of VAriance) followed by a post hoc Tukey’s HSD (Honest Significant Difference) test was performed;
3. for two-factor studies (e.g., Figures 3A and S2O’), two-way ANOVA was used;
4. for studies with more factors involved (e.g., Figure S1B), three-way ANOVA was used to determine whether and how three independent factors affect a dependent variable (e.g., serration number in Figure S1B);
5. for statistical validation of differences between genotypes (fixed factors or outer factors) with independent biological replicates (nested random factors or inner factors), nested ANOVA was used (e.g., Figures 1M and 1N), except for where different numbers of replicates were involved for each genotype (e.g., Figures 2M and 2N, non-parametric tests were used, with replicate information indicated in the plots.);
6. for studies involving covariate or categorical independent variables (e.g., leaf nodes in Figures 4G and 4J), the ANalysis of COVAriance (ANCOVA), a general linear model blending ANOVA and regression, was employed to test whether the means of the dependent variable (NDMC or leaflet number) were equal across levels of leaf nodes (covariate) and across different genotypes (categorical variable).

### o Non-parametric tests

For comparisons between two groups (e.g., Figure 4A), Welch’s *t*-test was used; for multiple comparisons (e.g., Figures 2M and 2N), the Kruskal-Wallis test was preformed, followed by post hoc Wilcoxon Rank Sum pairwise tests with Benjamini-Hochberg (BH or alias FDR) *p*-value adjustment.

### • Regressions

For scatter plots (e.g., temporal dynamics in Figure 3D or growth alignments in Figure 3F), regression is cubic polynomial (polynomial degree at 3), due to a good balance between the representativeness and simplicity of the fitting.^16^ For analysis of the heteroblastic changes (covariance) in leaf geometry (e.g., Figures 4G and 4J), linear models were constructed (*p* < 2.2E-16 for all the linear regressions), with coefficients (*R*^2^) and corresponding equations (intercepts and slopes) indicated.

The 95% confidence interval is shown in shade along these regression lines.

### • PCA

In leaf shape-space analysis, the first 64 harmonics of the Fourier coefficients were used for Principal Component Analysis (PCA). The first two components (PC1 and PC2) that account for the majority of shape variance were plotted along the *x*- and *y*-axis, respectively (Figures 1C and 2A). The values were normalized by the standard deviation of each principal component. The percentage of variance attributed to each component was indicated. Statistical features, such as mean (cross), standard error (ellipse), and distribution range (polygon) of each group, were plotted in the shape-space.

### • Hierarchical clustering

To assess the similarity between growth temporal dynamics (e.g., Figures 2D and S2G1-S2G4), hierarchical clustering trees were built using mean values of leaf-wide growth measures under different genetic backgrounds over time. Clustering was done using the Euclidean distance and complete lineage method. Heatmaps and relationship trees were visualized using Heatmapper.^59^

### Additional SI items

**Video S1** (MP4 file)

**Distal augmentation of *SPL9* balances adult leaf proximodistal histogenesis in *Arabidopsis thaliana*.** Cell lineages and proliferation are tracked in Col-0 and *pLMI1::SPL9r:Venus* leaf8s from 2 days after initiation (DAI). Related to Figure 1.

**Video S2** (MP4 file)

**Proximal expression of *SPL9* is sufficient to trigger leaf juvenile-adult transition in *Arabidopsis thaliana*.** The temporal dynamic of cell lineages and proliferation in Col-0, *pCUC2::SPL9r:Venus*, and *pRCO::SPL9r:Venus* leaf1s from 2 days after initiation (DAI). Related to Figure 2.

**Video S3** (MP4 file)

**Juvenile versus adult leaf growth patterns in *Cardamine hirsuta*.** Cellular growth is tracked in *C. hirsuta* juvenile (leaf1) versus adult (leaf8) leaf development (1-5 days after initiation, DAI). Cell lineages and proliferation are shown. Related to Figure 3.

## REFERENCES

1. Bhatia, N., Runions, A., and Tsiantis, M. (2021). Leaf Shape Diversity: From Genetic Modules to Computational Models. Annual Review of Plant Biology, Vol 72, 2021 72, 325–356. 10.1146/annurev-arplant-080720-101613.

2. Coen, E., Kennaway, R., and Whitewoods, C. (2017). On genes and form. Development 144, 4203–4213. 10.1242/dev.151910.

3. Hudson, A. (2000). Development of symmetry in plants. Annu Rev Plant Phys 51, 349–370. DOI 10.1146/annurev.arplant.51.1.349.

4. Klingenberg, C.P. (2010). Evolution and development of shape: integrating quantitative approaches. Nat Rev Genet 11, 623–635. 10.1038/nrg2829.

5. Palmer, A.R. (2004). Symmetry breaking and the evolution of development. Science 306, 828–833. 10.1126/science.1103707.

6. Carroll, S.B. (2008). Evo-devo and an expanding evolutionary synthesis: a genetic theory of morphological evolution. Cell 134, 25–36. 10.1016/j.cell.2008.06.030.

7. Grimes, D.T. (2019). Making and breaking symmetry in development, growth and disease. Development 146. 10.1242/dev.170985.

8. Ashby, E. (1948). Studies in the Morphogenesis of Leaves .1. An Essay on Leaf Shape. New Phytologist 47, 153–176. DOI 10.1111/j.1469-8137.1948.tb05098.x.

9. Goebel, K. (1908). Einleitung in die experimentelle Morphologie der Pflanzen (BG Teubner).

10. Poethig, R.S. (1990). Phase-Change and the Regulation of Shoot Morphogenesis in Plants. Science 250, 923–930. DOI 10.1126/science.250.4983.923.

11. Poethig, R.S., and Fouracre, J. (2024). Temporal regulation of vegetative phase change in plants. Dev Cell 59, 4–19. 10.1016/j.devcel.2023.11.010.

12. Li, X.M., Jenke, H., Strauss, S., Bazakos, C., Mosca, G., Lymbouridou, R., Kierzkowski, D., Neumann, U., Naik, P., Huijser, P., et al. (2024). Cell-cycle-linked growth reprogramming encodes developmental time into leaf morphogenesis. Current Biology 34, 541–556.e515. 10.1016/j.cub.2023.12.050.

13. Rodriguez, R.E., Schommer, C., and Palatnik, J.F. (2016). Control of cell proliferation by microRNAs in plants. Curr Opin Plant Biol 34, 68–76. 10.1016/j.pbi.2016.10.003.

14. Sarvepalli, K., Das Gupta, M., Challa, K.R., and Nath, U. (2019). Molecular cartography of leaf development - role of transcription factors. Curr Opin Plant Biol 47, 22–31. 10.1016/j.pbi.2018.08.002.

15. Satterlee, J.W., and Scanlon, M.J. (2019). Coordination of Leaf Development Across Developmental Axes. Plants (Basel) 8. 10.3390/plants8100433.

16. Zhang, Z., Runions, A., Mentink, R.A., Kierzkowski, D., Karady, M., Hashemi, B., Huijser, P., Strauss, S., Gan, X., Ljung, K., and Tsiantis, M. (2020). A WOX/Auxin Biosynthesis Module Controls Growth to Shape Leaf Form. Curr Biol 30, 4857–4868 e4856. 10.1016/j.cub.2020.09.037.

17. Kierzkowski, D., Runions, A., Vuolo, F., Strauss, S., Lymbouridou, R., Routier-Kierzkowska, A.L., Wilson-Sanchez, D., Jenke, H., Galinha, C., Mosca, G., et al. (2019). A Growth-Based Framework for Leaf Shape Development and Diversity. Cell 177, 1405–1418 e1417. 10.1016/j.cell.2019.05.011.

18. Schwab, R., Palatnik, J.F., Riester, M., Schommer, C., Schmid, M., and Weigel, D. (2005). Specific effects of microRNAs on the plant transcriptome. Dev Cell 8, 517–527. 10.1016/j.devcel.2005.01.018.

19. Rhoades, M.W., Reinhart, B.J., Lim, L.P., Burge, C.B., Bartel, B., and Bartel, D.P. (2002). Prediction of plant microRNA targets. Cell 110, 513–520. Doi 10.1016/S0092-8674(02)00863-2.

20. Vuolo, F., Mentink, R.A., Hajheidari, M., Bailey, C.D., Filatov, D.A., and Tsiantis, M. (2016). Coupled enhancer and coding sequence evolution of a homeobox gene shaped leaf diversity. Genes Dev 30, 2370–2375. 10.1101/gad.290684.116.

21. Wang, J.W., Czech, B., and Weigel, D. (2009). miR156-regulated SPL transcription factors define an endogenous flowering pathway in *Arabidopsis thaliana*. Cell 138, 738–749. 10.1016/j.cell.2009.06.014.

22. Wang, J.W., Schwab, R., Czech, B., Mica, E., and Weigel, D. (2008). Dual effects of miR156-targeted *SPL* genes and *CYP78A5/KLUH* on plastochron length and organ size in *Arabidopsis thaliana*. Plant Cell 20, 1231–1243. 10.1105/tpc.108.058180.

23. Wu, G., Park, M.Y., Conway, S.R., Wang, J.W., Weigel, D., and Poethig, R.S. (2009). The sequential action of miR156 and miR172 regulates developmental timing in *Arabidopsis*. Cell 138, 750–759. 10.1016/j.cell.2009.06.031.

24. Bilsborough, G.D., Runions, A., Barkoulas, M., Jenkins, H.W., Hasson, A., Galinha, C., Laufs, P., Hay, A., Prusinkiewicz, P., and Tsiantis, M. (2011). Model for the regulation of *Arabidopsis thaliana* leaf margin development. Proc Natl Acad Sci U S A 108, 3424–3429. 10.1073/pnas.1015162108.

25. Rubio-Somoza, I., Zhou, C.M., Confraria, A., Martinho, C., von Born, P., Baena-Gonzalez, E., Wang, J.W., and Weigel, D. (2014). Temporal control of leaf complexity by miRNA-regulated licensing of protein complexes. Curr Biol 24, 2714–2719. 10.1016/j.cub.2014.09.058.

26. Piazza, P., Bailey, C.D., Cartolano, M., Krieger, J., Cao, J., Ossowski, S., Schneeberger, K., He, F., de Meaux, J., Hall, N., et al. (2010). *Arabidopsis thaliana* leaf form evolved via loss of KNOX expression in leaves in association with a selective sweep. Curr Biol 20, 2223–2228. 10.1016/j.cub.2010.11.037.

27. Cartolano, M., Pieper, B., Lempe, J., Tattersall, A., Huijser, P., Tresch, A., Darrah, P.R., Hay, A., and Tsiantis, M. (2015). Heterochrony underpins natural variation in Cardamine hirsuta leaf form. Proc Natl Acad Sci U S A 112, 10539–10544. 10.1073/pnas.1419791112.

28. Bharathan, G., Goliber, T.E., Moore, C., Kessler, S., Pham, T., and Sinha, N.R. (2002). Homologies in leaf form inferred from *KNOXI* gene expression during development. Science 296, 1858–1860. 10.1126/science.1070343.

29. Hareven, D., Gutfinger, T., Parnis, A., Eshed, Y., and Lifschitz, E. (1996). The making of a compound leaf: genetic manipulation of leaf architecture in tomato. Cell 84, 735–744. 10.1016/s0092-8674(00)81051-x.

30. Hay, A., and Tsiantis, M. (2006). The genetic basis for differences in leaf form between *Arabidopsis thaliana* and its wild relative *Cardamine hirsuta*. Nat Genet 38, 942–947. 10.1038/ng1835.

31. Vlad, D., Kierzkowski, D., Rast, M.I., Vuolo, F., Dello Ioio, R., Galinha, C., Gan, X.C., Hajheidari, M., Hay, A., Smith, R.S., et al. (2014). Leaf Shape Evolution Through Duplication, Regulatory Diversification, and Loss of a Homeobox Gene. Science 343, 780–783. 10.1126/science.1248384.

32. Andres, R.J., Coneva, V., Frank, M.H., Tuttle, J.R., Samayoa, L.F., Han, S.W., Kaur, B., Zhu, L., Fang, H., Bowman, D.T., et al. (2017). Modifications to a *LATE MERISTEM IDENTITY1* gene are responsible for the major leaf shapes of Upland cotton (*Gossypium hirsutum* L.). Proc Natl Acad Sci U S A 114, E57–E66. 10.1073/pnas.1613593114.

33. Hu, L., Zhang, H., Sun, Y., Shen, X., Amoo, O., Wang, Y., Fan, C., and Zhou, Y. (2020). *BnA10.RCO*, a homeobox gene, positively regulates leaf lobe formation in *Brassica napus* L. Theor Appl Genet 133, 3333–3343. 10.1007/s00122-020-03672-3.

34. Sicard, A., Thamm, A., Marona, C., Lee, Young W., Wahl, V., Stinchcombe, John R., Wright, Stephen I., Kappel, C., and Lenhard, M. (2014). Repeated Evolutionary Changes of Leaf Morphology Caused by Mutations to a Homeobox Gene. Current Biology 24, 1880–1886. 10.1016/j.cub.2014.06.061.

35. Hay, A., and Tsiantis, M. (2010). KNOX genes: versatile regulators of plant development and diversity. Development 137, 3153–3165. 10.1242/dev.030049.

36. Tsuda, K., and Hake, S. (2015). Diverse functions of KNOX transcription factors in the diploid body plan of plants. Curr Opin Plant Biol 27, 91–96. 10.1016/j.pbi.2015.06.015.

37. Tsuda, K., and Hake, S. (2016). Homeobox Transcription Factors and the Regulation of Meristem Development and Maintenance. In Plant Transcription Factors, pp. 215–228. 10.1016/b978-0-12-800854-6.00014-2.

38. Rast-Somssich, M.I., Broholm, S., Jenkins, H., Canales, C., Vlad, D., Kwantes, M., Bilsborough, G., Dello Ioio, R., Ewing, R.M., Laufs, P., et al. (2015). Alternate wiring of a *KNOXI* genetic network underlies differences in leaf development of *A. thaliana* and *C. hirsuta*. Genes Dev 29, 2391–2404. 10.1101/gad.269050.115.

39. Wang, Y., Strauss, S., Liu, S., Pieper, B., Lymbouridou, R., Runions, A., and Tsiantis, M. (2022). The cellular basis for synergy between *RCO* and *KNOX1* homeobox genes in leaf shape diversity. Curr Biol 32, 3773–3784 e3775. 10.1016/j.cub.2022.08.020.

40. Bhatia, N., Wilson-Sánchez, D., Strauss, S., Vuolo, F., Pieper, B., Hu, Z.L., Rambaud-Lavigne, L., and Tsiantis, M. (2023). Interspersed expression of *CUP-SHAPED COTYLEDON2* and *REDUCED COMPLEXITY* shapes *Cardamine hirsuta* complex leaf form. Current Biology 33, 2977–2987. 10.1016/j.cub.2023.06.037.

41. Baumgarten, L., Pieper, B., Song, B., Mane, S., Lempe, J., Lamb, J., Cooke, E.L., Srivastava, R., Strutt, S., Zanko, D., et al. (2023). Pan-European study of genotypes and phenotypes in the Arabidopsis relative *Cardamine hirsuta* reveals how adaptation, demography, and development shape diversity patterns. PLoS Biol 21, e3002191. 10.1371/journal.pbio.3002191.

42. Chitwood, D.H., and Sinha, N.R. (2016). Evolutionary and Environmental Forces Sculpting Leaf Development. Curr Biol 26, R297–306. 10.1016/j.cub.2016.02.033.

43. Dennis, R.J., Whitewoods, C.D., and Harrison, C.J. (2019). Quantitative methods in like-for-like comparative analyses of Aphanorrhegma (Physcomitrella) patens phyllid development. Journal of Bryology 41, 314–321. 10.1080/03736687.2019.1668109.

44. Lawrence, E.H., Leichty, A.R., Doody, E.E., Ma, C., Strauss, S.H., and Poethig, R.S. (2021). Vegetative phase change in Populus tremula x alba. New Phytol 231, 351–364. 10.1111/nph.17316.

45. Leichty, A.R., and Poethig, R.S. (2019). Development and evolution of age-dependent defenses in ant-acacias. Proc Natl Acad Sci U S A 116, 15596–15601. 10.1073/pnas.1900644116.

46. Sederoff, R.R., Wang, J.-W., Park, M.Y., Wang, L.-J., Koo, Y., Chen, X.-Y., Weigel, D., and Poethig, R.S. (2011). MiRNA Control of Vegetative Phase Change in Trees. PLoS Genetics 7. 10.1371/journal.pgen.1002012.

47. Silva, P.O., Batista, D.S., Cavalcanti, J.H.F., Koehler, A.D., Vieira, L.M., Fernandes, A.M., Barrera-Rojas, C.H., Ribeiro, D.M., Nogueira, F.T.S., and Otoni, W.C. (2019). Leaf heteroblasty in Passiflora edulis as revealed by metabolic profiling and expression analyses of the microRNAs miR156 and miR172. Ann Bot 123, 1191–1203. 10.1093/aob/mcz025.

48. Struebe, S., Deibar, S., Rötzer, J., Mosiolek, M., Jandrasits, K., and Dolan, L. (2023). Meristem dormancy in *Marchantia polymorpha* is regulated by a liverwort-specific miRNA and a clade III *SPL* gene. Current Biology 33, 660-+. 10.1016/j.cub.2022.12.062.

49. Tsuzuki, M., Futagami, K., Shimamura, M., Inoue, C., Kunimoto, K., Oogami, T., Tomita, Y., Inoue, K., Kohchi, T., Yamaoka, S., et al. (2019). An Early Arising Role of the MicroRNA156/529-SPL Module in Reproductive Development Revealed by the Liverwort Marchantia polymorpha. Curr Biol 29, 3307–3314 e3305. 10.1016/j.cub.2019.07.084.

50. Costa, M.M.R., Yang, S., Critchley, J., Feng, X., Wilson, Y., Langlade, N., Copsey, L., and Hudson, A. (2012). The genetic basis for natural variation in heteroblasty in *Antirrhinum*. New Phytol 196, 1251–1259. 10.1111/j.1469-8137.2012.04347.x.

51. Doody, E., Zha, Y.Q., He, J., and Poethig, R.S. (2022). The genetic basis of natural variation in the timing of vegetative phase change in *Arabidopsis thaliana*. Development 149. 10.1242/dev.200321.

52. Clough, S.J., and Bent, A.F. (1998). Floral dip: a simplified method for *Agrobacterium*-mediated transformation of *Arabidopsis thaliana*. Plant J 16, 735–743. 10.1046/j.1365-313x.1998.00343.x.

53. Vuolo, F., Kierzkowski, D., Runions, A., Hajheidari, M., Mentink, R.A., Gupta, M.D., Zhang, Z., Vlad, D., Wang, Y., Pecinka, A., et al. (2018). LMI1 homeodomain protein regulates organ proportions by spatial modulation of endoreduplication. Genes Dev 32, 1361–1366. 10.1101/gad.318212.118.

54. Barbier de Reuille, P., Routier-Kierzkowska, A.L., Kierzkowski, D., Bassel, G.W., Schupbach, T., Tauriello, G., Bajpai, N., Strauss, S., Weber, A., Kiss, A., et al. (2015). MorphoGraphX: A platform for quantifying morphogenesis in 4D. Elife 4, 05864. 10.7554/eLife.05864.

55. Strauss, S., Runions, A., Lane, B., Eschweiler, D., Bajpai, N., Trozzi, N., Routier-Kierzkowska, A.L., Yoshida, S., Rodrigues da Silveira, S., Vijayan, A., et al. (2022). Using positional information to provide context for biological image analysis with MorphoGraphX 2.0. Elife 11. 10.7554/eLife.72601.

56. Sapala, A., Runions, A., Routier-Kierzkowska, A.L., Das Gupta, M., Hong, L., Hofhuis, H., Verger, S., Mosca, G., Li, C.B., Hay, A., et al. (2018). Why plants make puzzle cells, and how their shape emerges. Elife 7. 10.7554/elife.32794.

57. Mary, H., and Brouhard, G.J. (2019). Kappa (*κ*): Analysis of Curvature in Biological Image Data using B-splines. bioRxiv, 852772. 10.1101/852772.

58. Leigh, A., Sevanto, S., Close, J.D., and Nicotra, A.B. (2016). The influence of leaf size and shape on leaf thermal dynamics: does theory hold up under natural conditions? Plant, Cell & Environment 40, 237–248. 10.1111/pce.12857.

59. Babicki, S., Arndt, D., Marcu, A., Liang, Y., Grant, J.R., Maciejewski, A., and Wishart, D.S. (2016). Heatmapper: web-enabled heat mapping for all. Nucleic Acids Res 44, W147–153. 10.1093/nar/gkw419.

