## Supplemental info_Fig.S1-S4, table S1 for "Age-associated growth control modifies leaf proximodistal symmetry and enables leaf shape diversification"

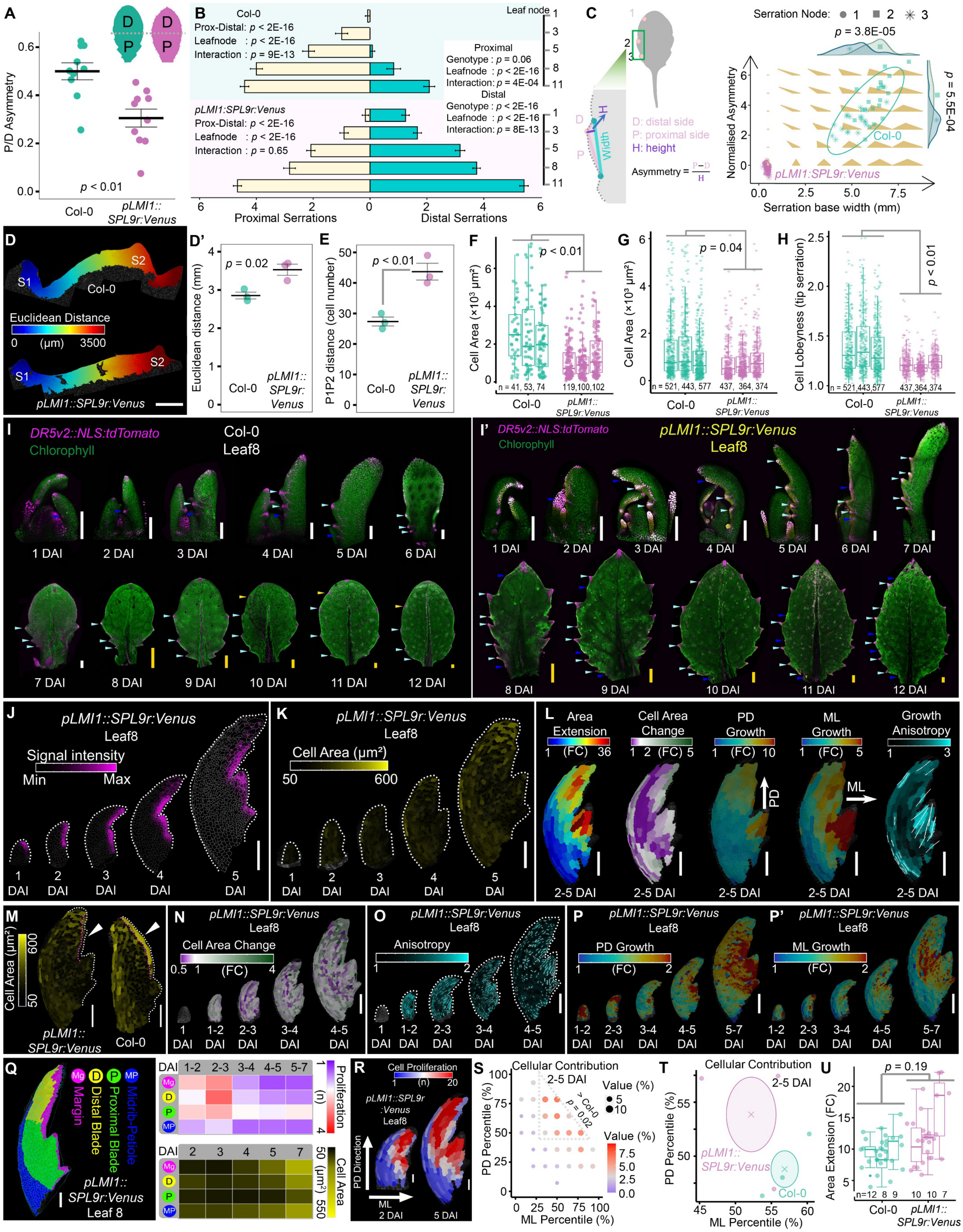
A
P/D Asymmetry
p < 0.01
Col-0
pLMI1::SPL9r:Venus
B
Col-0
Prox-Distal: p < 2E-16
Leafnode : p < 2E-16
Interaction : p = 9E-13
Leaf node
Proximal
Genotype : p = 0.06
Leafnode : p < 2E-16
Interaction: p = 4E-04
Distal
Genotype : p < 2E-16
Leafnode : p < 2E-16
Interaction: p = 8E-13
pLMI1:SPL9r:Venus
Prox-Distal: p < 2E-16
Leafnode : p < 2E-16
Interaction : p = 0.65
Proximal Serrations
Distal Serrations
C
D: distal side
P: proximal side
H: height
Width
Asymmetry = (P−D)/H
Serration Node:
p = 3.8E-05
p = 5.5E-04
Normalised Asymmetry
Col-0
pLMI1:SPL9r:Venus
Serration base width (mm)
D
S1
S2
Col-0
Euclidean Distance
0 (μm) 3500
pLMI1::SPL9r:Venus
D'
p = 0.02
Euclidean distance (mm)
E
p < 0.01
P1P2 distance (cell number)
F
p < 0.01
Cell Area (×10³ μm²)
n = 41, 53, 74
119,100,102
G
p = 0.04
n = 521, 443, 577
437, 364, 374
H
p < 0.01
Cell Lobeyness (tip serration)
I
DR5v2::NLS:tdTomato
Chlorophyll
Col-0
Leaf8
1 DAI
2 DAI
3 DAI
4 DAI
5 DAI
6 DAI
7 DAI
8 DAI
9 DAI
10 DAI
11 DAI
12 DAI
I'
DR5v2::NLS:tdTomato
Chlorophyll
pLMI1::SPL9r:Venus
Leaf8
J
pLMI1::SPL9r:Venus
Leaf8
Signal intensity
Min
Max
K
pLMI1::SPL9r:Venus
Leaf8
Cell Area (μm²)
50
600
L
Area Extension
Cell Area Change
PD Growth
ML Growth
Growth Anisotropy
(FC)
PD
ML
2-5 DAI
M
Cell Area (μm²)
pLMI1::SPL9r:Venus
Col-0
N
pLMI1::SPL9r:Venus
Leaf8
Cell Area Change
(FC)
O
pLMI1::SPL9r:Venus
Leaf8
Anisotropy
P
pLMI1::SPL9r:Venus
Leaf8
PD Growth
(FC)
P'
pLMI1::SPL9r:Venus
Leaf8
ML Growth
(FC)
Q
Mg Margin
D Distal Blade
P Proximal Blade
MP Midrib-Petiole
pLMI1::SPL9r:Venus
Leaf 8
DAI 1-2 2-3 3-4 4-5 5-7
Proliferation (n)
DAI 2 3 4 5 7
Cell Area (μm²)
R
Cell Proliferation
(n)
pLMI1::SPL9r:Venus
Leaf8
PD Direction
ML
2 DAI
5 DAI
S
Cellular Contribution
2-5 DAI
Value (%)
> Col-0
p = 0.02
PD Percentile (%)
ML Percentile (%)
T
Cellular Contribution
2-5 DAI
pLMI1::SPL9r:Venus
Col-0
PD Percentile (%)
ML Percentile (%)
U
p = 0.19
Area Extension (FC)
n=12 8 9
10 10 7
Col-0
pLMI1::SPL9r:Venus

**Figure S1. Distal augmentation of *SPL9* is sufficient to change proximodistal asymmetry in *Arabidopsis thaliana* adult leaves, Related to Figure 1**

(A) Proximodistal asymmetry of leaf blade outline. P, proximal; D, distal. Asymmetry was calculated using the curvature of the proximal versus distal blade (see STAR Methods). Adult leaves (leaf10) from wild-type and *pLMI1::SPL9r:Venus* background are compared. Mean ± SE; n = 10. *p*, Student's *t*-test. Representative samples are shown on the top with color key.
(B) *pLMI1::SPL9r:Venus* potentiates serration formation, especially in leaf distal parts. Serration distribution was surveyed in *pLMI1::SPL9r:Venus* and Col-0 mature leaves. n = 12 independent replicates. A three-way ANOVA showed that genotype, leaf node, and leaf zone (distal versus proximal parts) all have strong effects on serration number ($p$ < 3E-10). Serrations (the response variable) along proximal versus distal margins (a categorical variable) of different leaf nodes (another independent categorical variable) were then compared with two-way ANOVA in Col-0 or *pLMI1::SPL9r:Venus,* separately. Furthermore, proximal or distal serrations (continuous dependent variable) in different leaf nodes (one categorical variable) under different genetic backgrounds (the second categorical variable) were compared. Note that the proximal distribution of serrations is similar in both genotypes, whereas the *SPL9* distal augmentation under the *LMI1* promoter (over-) rescues serration development in distal blades.
(C) Serration symmetry analysis in mature leaf10s. Serrations were numbered from the leaf tip, following their order of biological development, with the tip serration excluded. Asymmetry was defined as the difference between proximal and distal sides, and normalized by height. Serrations in *pLMI1::SPL9r:Venus* and Col-0 were measured and plotted as serration bilateral asymmetry versus its size (base width). Background brown triangles, morphospace for serration variations. Ellipse, 95% confidence interval. n = 20 serrations from ten leaves (both sides). The first serration fades over time in Col-0 fully expanded leaf10, thus is unmeasurable. The differences in asymmetry (distribution plot on the right) and width (top plot) between the second and the third serrations in Col-0 were tested by Student's *t*-test; there are no significant differences in either measure among *pLMI1::SPL9r:Venus* serrations.
(D-F) Serration spacing. The distance between the first two proximal serrations in leaf8 was measured in Euclidean distance (D, heatmaps of representatives; D', quantification) and cell number (E, see Figure 1E for representative heatmaps). The areas of marginal cells between these two serrations was also compared (F). Mean ± SE, n = 3. *p*, Student's *t*-test.
(G and H) Serration maturation (indicated by cell area and lobeyness). Tip serrations (a radius of 600 µm from the tip cell) from mature leaves were examined. n = 3 leaf8 replicates for each genotype; numbers of cells plotted for each replicate (box plot) are indicated. *p*-value, nested ANOVA.
(I and I') Serration establishment in Col-0 (n > 6) and *pLMI1::SPL9r:Venus* (n > 10) adult leaves. Representative leaf8 samples at different stages are shown, with developing serrations marked by *DR5v2::NLS:tdTomato* (auxin maxima, magenta). Triangles indicate serrations on one side: cyan triangles, prominent serrations; blue, emerging serrations; yellow, fading serrations. Scale bars: 100 µm for 1-7 DAI in white; 500 µm, 8-12 DAI, yellow.
(J) *pLMI1::SPL9r:Venus* is predominantly expressed in the distal margin in leaf development. Heatmaps show the epidermal projection (2-6 µm depth) of Venus signal in a representative sample. n = 4. Scale bar, 100 µm.
(K-P') Heatmaps of cell area (K) and its change (N), growth anisotropy (O), and proximodistal (PD) versus mediolateral (ML) growth (P and P') during *pLMI1::SPL9r:Venus* leaf8 development. Measures were projected on the later time points of the indicated periods in (N and O) while on the earlier time points in (P and P'). Growth from 2-5 DAI was summarized on 5 DAI meshes (L), and the significant difference in distal margin differentiation between *pLMI1::SPL9r:Venus* and Col-0 is highlighted by cell area at 5 DAI (M). White lines on anisotropy heatmaps indicate maximum Principal Directions of Growth (PDG)[S1] in cells with high anisotropy: > 2 (L); or > 1.2 (O). FC, fold change. n = 4 replicates. Scale bars, 100 µm.
(Q) Temporal changes in proliferation and cell area in different cell types. Cell types with distinct developmental features were defined at the end of the observation period (7 DAI) and were tracked backward based on developmental lineages. Mg, margin; D, distal blade; P, proximal blade; MP, midrib-petiole. The average growth of each cell type is visualized as heat values across the observation period. n = 4.
(R-T) Spatial redistribution of cell proliferation and consequent cellular contribution from 2-5 DAI in response to *pLMI1::SPL9r:Venus.* (R), proliferation from 2-5 DAI is projected on both 2 DAI (with PD-ML coordinate axes indicated; scale bar, 20 µm) and 5 DAI meshes (scale bar, 50 µm). Cellular contribution from 2-5 DAI in *pLMI1::SPL9r:Venus* leaf8 was plotted along the PD and ML directions (S). The distal cells (in the dashed triangle) at 2 DAI contribute more to later cell populations in *pLMI1::SPL9r:Venus* than in wild-type (Col-0) leaf8[S2]. *p*-value, two-way ANOVA. In this way, *pLMI1::SPL9r:Venus* induces an acropetal shift of the weighted mean of cellular contribution, compared to wild-type (T). Ellipse, defined by the standard deviation in both directions; cross, mean. The wild-type samples from a previous study[S2] were reanalyzed as controls. Three replicates were used for each genotype.
(U) Equivalent areal growth of distal margins (*LMI1* domain) in Col-0 and *pLMI1::SPL9r:Venus* leaf8. *p*-value, nested ANOVA.

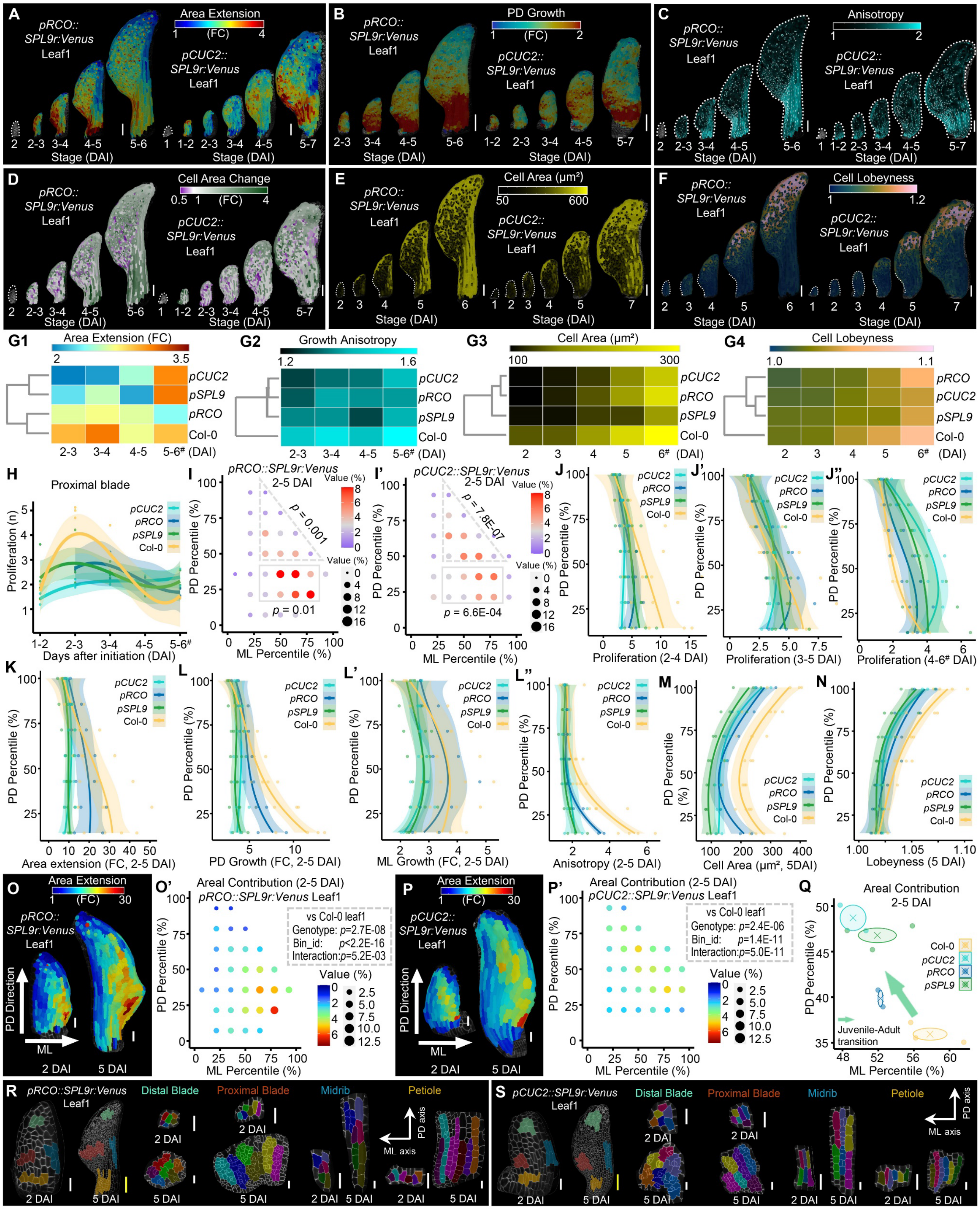

**Figure S2. Cellular growth reprogramming upon proximal expression of *SPL9* in *Arabidopsis thaliana* leaf1, Related to Figure 2**

(A-F) Heatmaps of area extension (A), growth along the proximodistal axis (PD growth, B), anisotropy (C), cell area change (D), cell area (E), and cell lobeyness (F) in leaf1 with SPL9 proximal supplementation (*pRCO::SPL9r:Venus* and *pCUC2::SPL9r:Venus*). Measures over two consecutive days are visualized on the earlier (B) or later meshes (A, C, and D). DAI, days after initiation. Indistinct parts of primordia are outlined with dotted lines. White lines in (C) indicate the growth direction where anisotropy > 1.2. Scale bar, 100 µm.
(G1-G4) Temporal patterns of cellular growth in leaf1 samples with disturbed *SPL9* expression. Measures of leaf-wide growth (mean), including area extension (G1), anisotropy (G2), cell area (G3), and cell lobeyness (G4), were clustered using the Euclidean distance complete lineage method. FC, fold change.
(H-J") Spatiotemporal reprogramming of cell proliferation upon *SPL9* manipulations in leaf1. (H), proliferation in proximal blades was tracked during development of indicated leaves, showing the elimination of the "proliferation burst" by ectopic SPL9. (I and I'), spatial redistribution of cellular contribution by *pRCO::SPL9r:Venus* and *pCUC2::SPL9r:Venus* during leaf1 early development (2-5 DAI) as examined by 2D (PD and ML directions) alignment. The contributions of cells in the proximal (indicated in rectangles) and distal (in dashed triangles) parts of blades were compared to wild-type (Col-0) leaf1[S2]. *p*-values, two-way ANOVA. Note that the contribution of proximal cells to later leaf development is reduced while that of distal cells is increased in both lines compared with Col-0 leaf1, leading to an acropetal shift of the overall cellular contribution (see Figure 2E). (J-J"), one-dimensional (PD direction) alignment of proliferation was established at different developmental stages (J, early-stage, 2-4 DAI; J', mid-stage, 3-5 DAI; J", late-stage, 4-6 DAI), showing temporal changes in cell proliferation and its spatial distribution along the PD axis.
(K-N) PD alignment of cellular growth, including area extension (K), growth in the PD and ML directions (L and L'), growth anisotropy (L"), cell area (M), and lobeyness (N). Growth from 2-5 DAI was aligned to the PD axes of 2 DAI primordia. Note that PD growth and anisotropy are significantly repressed at primordia bases by ectopic *SPL9*, in association with suppressed petiole elongation.
(O-Q) Areal growth changes upon local (proximal) SPL9 action. (O and P), area extension from 2-5 DAI is projected on both 2 DAI (with PD-ML coordinates illustrated) and 5 DAI in *pRCO::SPL9r:Venus* and *pCUC2::SPL9r:Venus*. Scale bar, 20 µm for 2 DAI, 50 µm for 5 DAI samples. (O' and P'), the contributions of 2 DAI cells to 5 DAI leaf areas were aligned in 2D coordinates and compared to the wild-type Col-0 leaf1 pattern[S2]. A significant diminution and apparent acropetal redistribution of growth contributions upon SPL9 modulation was found in both lines (two-way ANOVA). (Q), the spatial redistribution (weighted means) of areal contribution from 2-5 DAI in leaf1 by *SPL9* misexpression is summarized. Ellipse, the standard deviation of the mean (inner cross) along PD and ML axes.
(R and S) Cellular growth tracking of representative lineages in divergent developmental domains. During tracking, different colors identify sectors or cells. Note the enhanced cell proliferation and concomitantly delayed cell maturation in proximal blades, as well as the repressed elongation of petiole sectors in both lines compared to Col-0 leaf1[S2]. Scale bars: 20 µm, except for the yellow bars for 5 DAI primordia (100 µm).
Fitting lines in (H and J-N) is cubic polynomial regression with a 95% confidence interval (shaded). Genotypes in (G1-N) are given in short: *pRCO*, *pRCO::SPL9r:Venus* (n = 2); *pCUC2*, *pCUC2::SPL9r:Venus* (n = 2). *pSPL9*, *pSPL9::SPL9r:Venus* (n = 3) and Col-0 (n = 4) from Li et al.[S2] were reassessed as positive and background controls, respectively. [#], 7 DAI for *pCUC2::SPL9r:Venus* and *pSPL9::SPL9r:Venus.*

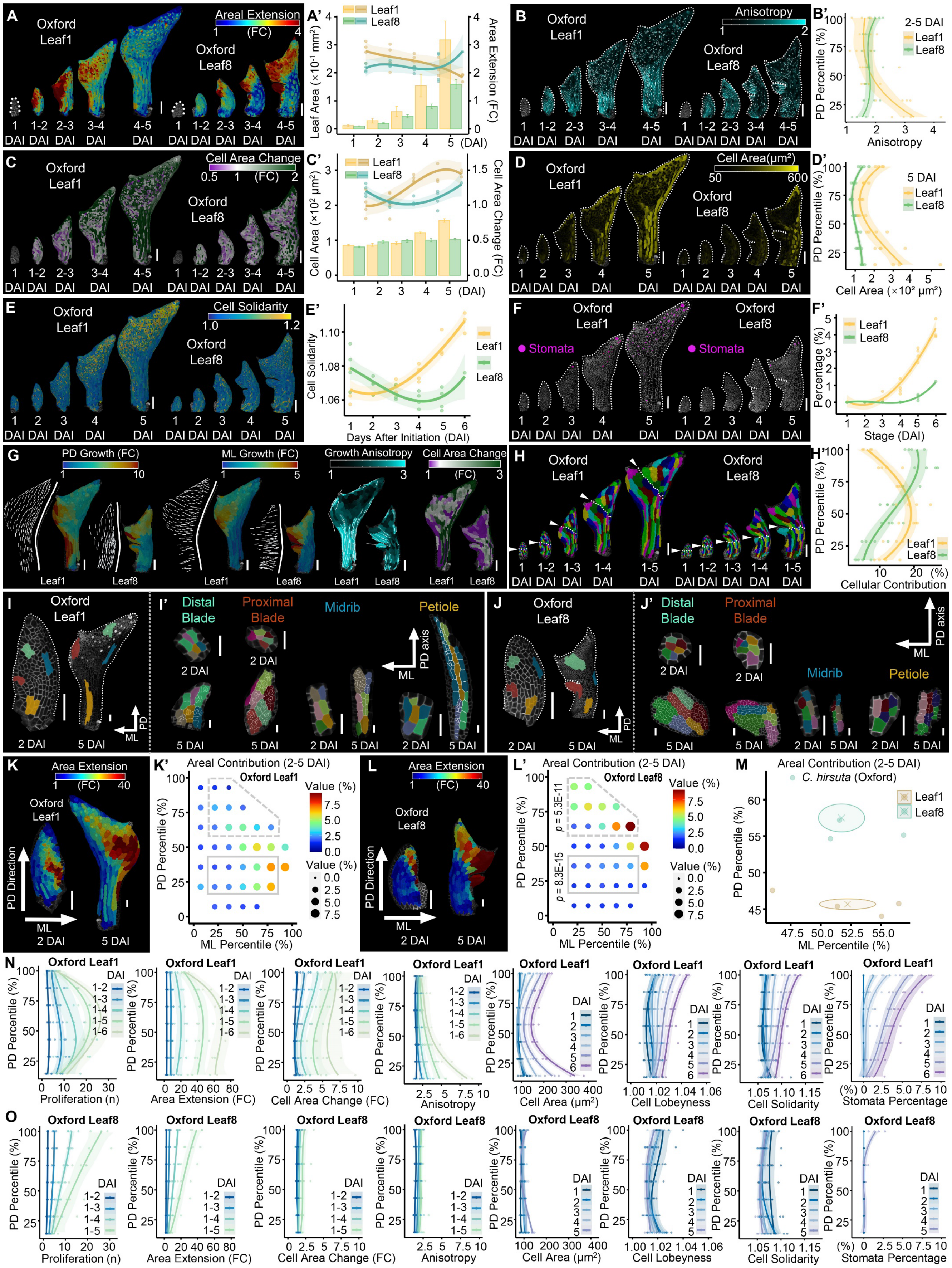

A
Oxford Leaf1
Oxford Leaf8
Areal Extension
1 (FC) 4
1 DAI
1-2 DAI
2-3 DAI
3-4 DAI
4-5 DAI
A'
Leaf1
Leaf8
Leaf Area (×10⁻¹ mm²)
Area Extension (FC)
B
Anisotropy
B'
2-5 DAI
PD Percentile (%)
Anisotropy
C
Cell Area Change
0.5 1 (FC) 2
C'
Cell Area (×10² μm²)
Cell Area Change (FC)
D
Cell Area(μm²)
50 600
D'
5 DAI
Cell Area (×10² μm²)
E
Cell Solidarity
1.0 1.2
E'
Cell Solidarity
Days After Initiation (DAI)
F
Stomata
F'
Percentage (%)
Stage (DAI)
G
PD Growth (FC)
ML Growth (FC)
Growth Anisotropy
Cell Area Change
H
1-3 DAI
1-4 DAI
1-5 DAI
H'
Cellular Contribution
I
I'
Distal Blade
Proximal Blade
Midrib
Petiole
PD axis
ML
2 DAI
5 DAI
J
J'
K
Area Extension
1 (FC) 40
PD Direction
K'
Areal Contribution (2-5 DAI)
Value (%)
ML Percentile (%)
L
L'
p = 5.3E-11
p = 8.3E-15
M
C. hirsuta (Oxford)
N
Proliferation (n)
Area Extension (FC)
Cell Area Change (FC)
Cell Area (μm²)
Cell Lobeyness
Cell Solidarity
Stomata Percentage
O

**Figure S3. Cellular growth framework for *Cardamine hirsuta* heteroblastic transition, Related to Figure 3**

(A-F) Spatiotemporal patterns of cellular growth (A and A', area extension; B and B', anisotropy; C and C', cell area change) and maturation (D and D', cell area; E and E', cell solidarity; and F and F', stomata emergence). Representative heatmaps are shown with indistinct primordia outlined. Cellular growth over one day was projected on the latter mesh (A-C). White lines in (B), maximum Principal Directions of Growth (PDG[S1] growth direction) of cells with high anisotropy (>1.2). Temporal dynamics of area extension (A') and cell area change (C') (right axes, scatter plots with fitting lines) were plotted together with consequent changes in leaf area (A') and mean cell area (C') (left *y*-axes; bars, mean ± SE), respectively. To show the spatial distribution, anisotropy (B', 2-5 DAI) and cell area (D', 5 DAI) were plotted as a function of the distance from the leaf base. Leaf-wide average of cell solidarity (E'), and stomata percentage (F') were tracked over time with trend lines. Note, cell expansion (C-D') and differentiation (E-F') are remarkably retarded in leaf8 compared to leaf1, highlighting the differentiation delay as an evolutionarily conserved feature in heteroblastic reprogramming in both *C. hirsuta* and *A. thaliana*[S2].

(G) Divergent growth features in *C. hirsuta* leaf1 versus leaf8 early development (2-5 DAI). Heatmaps of growth (area extension in fold change, FC) along the proximodistal (PD) and mediolateral (ML) axes (directions indicated by white lines) of developing leaves are visualized on representative samples. Growth anisotropy and cell area change from 2-5 DAI are visualized on 5 DAI meshes as well. White lines on anisotropy heatmaps indicate growth orientations (maximum PDG) when anisotropy > 2. These data highlight the premature differentiation (accelerated elongation along the proximodistal axis) of the petiole in leaf1 compared to leaf8, which resembles the divergent growth patterns in *A. thaliana* juvenile versus adult leaves.[S2]

(H and H') Cellular fate-mapping of *C. hirsuta* juvenile (leaf1 without leaflet) versus adult (leaf8, with leaflet emerging) leaf early development (1-5 DAI). (H), cellular lineages. Cells at 1 DAI are identified by different colors, and these identities were tracked in one-day time windows until 5 DAI. The developmental contribution of the proximal versus distal half of 1 DAI primordium is highlighted in both leaves (triangles and dashed lines track the PD midlines of 1 DAI primordia). (H'), quantification of cellular contribution of 2 DAI cells to 5 DAI primordia, as a function of distance from base. These results show that in leaf1, the proximal parts of primordia contribute significantly more than the distal parts to later development, whereas in leaf8, both parts make equivalent contributions until 5 DAI. This unbalanced PD developmental contribution in leaf1 compared to leaf8, as also seen in *A. thaliana*,[S2] indicates juvenile patterns are characterized by an accelerated basipetal cell differentiation or tissue maturation in both *C. hirsuta* and *A. thaliana*[S2].

(I-J') Growth tracing of divergent cell groups (in different colors; I, leaf1; J, leaf8) from 2-5 DAI. (I' and J'), cellular-scale close-ups of sectors. Colors reflect sector (I and J) or cell (I' and J') identity.

(K-M) Distribution of areal growth and its developmental contribution during *C. hirsuta* leaf1 and leaf8 early development (2-5 DAI). Cellular area extension from 2-5 DAI was projected on both 2 DAI and 5 DAI meshes of Oxford leaf1 (K) and leaf8 (L). The organ (PD/ML) coordinates were established at 2 DAI, and primordia were binned into seven equivalent groups along each direction (see STAR Methods). The contribution of each bin to the area of 5 DAI leaves was computed and normalized by the area of the starting bin to remove any binning effect. The mean cellular contributions of each bin were plotted (K' and L'). The areal contribution in leaf1 compared to leaf8 is smaller for distal cells (indicated in dashed polygons), while bigger for proximal cells (in grey rectangles), which echoes observations in *A. thaliana*.[S2] *p*-values, two-way ANOVA. The spatial distributions of these areal contributions (weighted centers) in both leaf nodes were summarized in the PD-ML coordinates (M). Cross, mean; ellipse, standard deviations along PD and ML directions. See Figure 3G-3K for the corresponding patterns of cellular contribution.

(N and O) PD alignment of daily growth in *C. hirsuta* leaf1 (N) versus leaf8 (O). Growth in increasing periods from 1 DAI was measured and aligned to the PD axis of the start primordium (1 DAI), while cell features (area, lobeyness, and solidarity) and stomata were aligned along the PD axes of the measured primordia. These measures indicate sustained proliferative growth and accelerated maturation at cellular and tissue levels in leaf1 compared to leaf8, which mimics the precocious development of leaf1 versus leaf8 in *A. thaliana*.[S2]

Four biological replicates were used for both leaf1 and leaf8. Fitting lines (A'-F', H', N and O), cubic regression with 95% confidence interval (shaded). Scale bars: 100 μm in (A-H); 50 μm (I, J, K, and L); 20 μm (I' and J'). DAI, days after initiation.

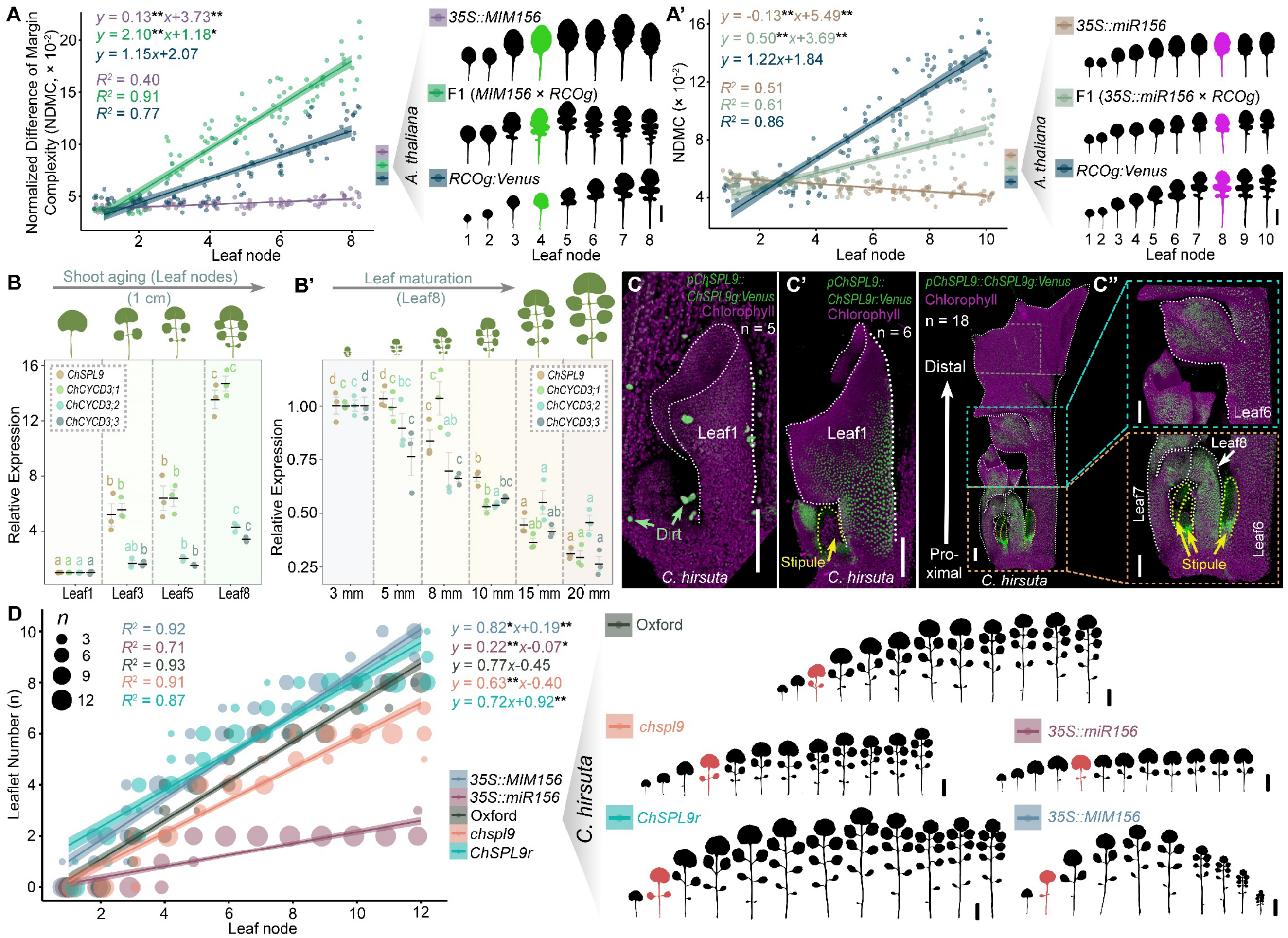


**Figure S4. Age-dependent growth reprogramming by miR156-SPL9 as an enabler of leaf complexity. Related to Figure 4**

(A and A') miR156 empowers RCO to sculpt leaf lobes in an age-dependent manner in *Arabidopsis thaliana*. Leaf lobe emergence was quantified by a Normalized Difference of Margin Complexity index (NDMC).[S3] *miR156* knockdown (*35S::MIM156*) enhances RCO's ability to create leaf lobes (A), whereas its overexpression (*35S::miR156*) imperils this ability (A'). Linear regressions indicate positive correlations (coefficient of determination, $R^2$) between NDMC and leaf nodes (shoot age). Accelerated (or enhanced, by *35S::MIM156*) or retarded (reduced, by *35S::miR156*) lobe formation upon RCO action was tested by comparing the indicated combination to *RCOg:Venus* using ANCOVA (both slope and intercept were tested). *, $p < 0.05$; **, $p < 0.01$. n = 10. Representative leaf series of each genotype are shown, with leaf4 (A) and leaf8 (A') colored in green and magenta, respectively, to highlight differences among genotypes. Scale bar, 1 cm.

(B and B') RT-qPCR assays showing the temporal expression pattern of the *SPL9-CYCD3* module during development of complex leaves in *Cardamine hirsuta*. Changes in gene expressions were examined over the progression of shoot aging (B, leaves at different nodes with the same length) and leaf maturation (B', leaf8 samples at different stages). Lowercase letters indicate the statistical significance for each gene (grouped by color) from Tukey's Honest Significant Difference (HSD) after one-way ANOVA. Error bars, Mean ± SE, n = 3.

(C-C'') Age-dependent expression of *ChSPL9* under the control of miR156. The native *ChSPL9* gene is expressed in adult leaves (C'') and not juvenile leaves (C, fluorescent dirt particles prove the actual presence of excitation and fluorescence detection). This temporal pattern is regulated by miR156, because the miR156-resistant allele (*ChSPL9r*) shows high expression in leaf1 (C'). During leaf maturation, *ChSPL9* expression decreases from distal (terminal leaflet) to proximal parts of leaves (basipetal decline), while remaining at a high level at the bases of leaflets (C'', see also Figure 4I). Five tiles were stitched for this image (C'', separation lines indicated), with the proximal two tiles highlighted in close-ups. Scale bars, 100 μm.

(D) The miR156-SPL9 pathway is necessary and sufficient for an age-dependent increase of leaflet number (heteroblasty) in *C. hirsuta*. *ChSPL9r,* short for *pChSPL9::ChSPL9r:Venus*, a miR156-resistant version of *ChSPL9* tagged with Venus-coding sequence. Linear regressions (fitting lines with 95% confidence interval, shaded) indicate a significant correlation ($R^2$ provided to indicate goodness-of-fit) between leaflet number and leaf node (shoot age). ANCOVA test was applied to compare the linear model of the indicated genotype to that of wild-type Oxford. Both slope (heteroblastic rate) and intercept were tested: *, $p < 0.05$; **, $p < 0.01$. Representative leaf silhouettes for each genotype are shown with the legend. The first leaf that generates a pair of leaflets in each leaf series is highlighted in dark red, as a marker of a shift in heteroblasty. Scale bars, 2 cm.

| Primer name | Oligos (5'-3') | Usage |
|---|---|---|
| NotIXmaI-SPL9-F[S2] | ATATATGgcggccgccccgggATGGAGATGGGTTCCAACTCGGGTC | Construction |
| NotIXmaI-Venus-R[S2] | ATATATgcggccgcCcccgggTCAATTGTACAGCTCGTCCATGCC | Construction |
| LB-1.3[S4] | ATTTTGCCGATTTCGGAAC | Genotyping |
| spl15-LP[S2] | TGTTGGTGTCTGAAGTTGCTG | Genotyping |
| spl15-R[S2] | AGGAAGCCAAAACCATAATGG | Genotyping |
| spl9-dCAPS-F1[S2] | TTCCTCCACTGAGTCATCCTC | Genotyping |
| spl9-dCAPS-R2[S2] | CCATCCCACAACCTTCCACTTGGCACCTTGaTAT | Genotyping |
| pSPL9-L[S2] | ACCTGATCGATCTCCTAACC | Genotyping |
| pMDC32-U1[S2] | GCTCTAGAACTAGTGGATCC | Genotyping |
| miR156site-F[S2] | GTGCTCTCTCTCTTCTGTCA | Genotyping |
| VENUS-seq[S2] | CTTGTACAGCTCGTCCATGCC | Genotyping |
| pSPL9-U[S2] | TCCCTCGTTTTGCTATGTGG | Genotyping |
| miR156r-R[S2] | GCTTAACAAGCTCAATGCGC | Genotyping |
| 35S-U[S2] | GTGGATTGATGTGACATCTC | Genotyping |
| gSPL9-L2[S2] | GAAAGAGGATGACTCAGTGG | Genotyping |
| miR156r-U[S2] | GCGCATTGAGCTTGTTAAGC | Genotyping |
| Venus-L2[S2] | GTGGTGCAGATCAGCTTCAG | Genotyping |
| mTFP1-F3[S2] | CTATCACTTTGTGGACCACC | Genotyping |
| SPL9genotyp-r[S2] | AACCTTCCACTTGGCACCTTGGTATC | Genotyping |
| pLMI1-U[S2] | AAGGTACTGTTGGGTTGCTC | Genotyping |
| pRCO-U | TAGCTCTAACGCTCGTATGG | Genotyping |
| pCUC2-U | CCCTTACTCAAGAACCATCC | Genotyping |
| pChSPL9-U1 | GTCTATTACCACTCTCGTCTC | Genotyping |
| as2-163_gt_F[S5] | GAGCTTCACCCTTCACAACGTGAAGCTG | Genotyping |
| as2-163_gt_R[S5] | GCTGACGAAGCTGATGTTGGAG | Genotyping |
| chspl9_F[a] | CCCACAACAGTCAAAATCAGG | Genotyping |
| chspl9_R[a] | CACCTCCAGCGTCTTCAAAG | Genotyping |
| ChSPL9-sgRNA1[S6] | CCGGGTCAGGCAGAGTCCGG | CRISPR-Cas9 |
| ChSPL9-sgRNA2[S6] | TCAAACAGACGGGTCCGTGG | CRISPR-Cas9 |
| ChSPL9-U | CAAGGTTCAGTTGGTGGAGGA | qPCR |
| ChSPL9-L | AGCCATTGTAACTGTCATGGG | qPCR |
| ChCYCD31-U | GTTATCTCCGATTCGAGATTTG | qPCR |
| ChCYCD31-L | TTGGATCTGTAAACCGATGCG | qPCR |
| ChCYCD32-U | CTTAACCCAAGCAAGAAGAGG | qPCR |
| ChCYCD32-L | GAGGACTAGTGATCACATCG | qPCR |
| ChCYCD33-U | CCAATCGGTGTGTTCGATGC | qPCR |
| ChCYCD33-L | CTACACGCCGAGAAACATTC | qPCR |
| ChGAPDH-F[S7] | TGACCACCGTCCACTCCATCAC | qPCR |
| ChGAPDH-R[S7] | GCTCTTCCACCTCTCCAGTCCTTC | qPCR |

**Table S1. Oligonucleotides used in the study, Related to STAR Methods.**

[a], CAPS (cleaved amplified polymorphic sequences) marker, *Bsa*W I digestion, *chspl9-1* and *chspl9-2*: 278 bp; wild-type (Oxford): 194+84 bp.

### Supplemental references


S1. Barbier de Reuille, P., Routier-Kierzkowska, A.L., Kierzkowski, D., Bassel, G.W., Schupbach, T., Tauriello, G., Bajpai, N., Strauss, S., Weber, A., Kiss, A., et al. (2015). MorphoGraphX: A platform for quantifying morphogenesis in 4D. Elife *4*, 05864. 10.7554/eLife.05864.

S2. Li, X.M., Jenke, H., Strauss, S., Bazakos, C., Mosca, G., Lymbouridou, R., Kierzkowski, D., Neumann, U., Naik, P., Huijser, P., et al. (2024). Cell-cycle-linked growth reprogramming encodes developmental time into leaf morphogenesis. Current Biology *34*, 541-556.e515. 10.1016/j.cub.2023.12.050.

S3. Leigh, A., Sevanto, S., Close, J.D., and Nicotra, A.B. (2016). The influence of leaf size and shape on leaf thermal dynamics: does theory hold up under natural conditions? Plant, Cell & Environment *40*, 237-248. 10.1111/pce.12857.

S4. O'Malley, R.C., Barragan, C.C., and Ecker, J.R. (2015). A user's guide to the Arabidopsis T-DNA insertion mutant collections. Methods Mol Biol *1284*, 323-342. 10.1007/978-1-4939-2444-8_16.

S5. Wang, Y., Strauss, S., Liu, S., Pieper, B., Lymbouridou, R., Runions, A., and Tsiantis, M. (2022). The cellular basis for synergy between *RCO* and *KNOX1* homeobox genes in leaf shape diversity. Current Biology *32*, 3773-3784 e3775. 10.1016/j.cub.2022.08.020.

S6. Baumgarten, L., Pieper, B., Song, B., Mane, S., Lempe, J., Lamb, J., Cooke, E.L., Srivastava, R., Strutt, S., Zanko, D., et al. (2023). Pan-European study of genotypes and phenotypes in the Arabidopsis relative *Cardamine hirsuta* reveals how adaptation, demography, and development shape diversity patterns. PLoS Biology *21*, e3002191. 10.1371/journal.pbio.3002191.

S7. Vlad, D., Kierzkowski, D., Rast, M.I., Vuolo, F., Dello Ioio, R., Galinha, C., Gan, X.C., Hajheidari, M., Hay, A., Smith, R.S., et al. (2014). Leaf Shape Evolution Through Duplication, Regulatory Diversification, and Loss of a Homeobox Gene. Science *343*, 780-783. 10.1126/science.1248384.
